# A CRISPR-induced DNA break can trigger crossover, chromosomal loss and chromothripsis-like rearrangements

**DOI:** 10.1101/2023.05.22.541757

**Authors:** Aviva Samach, Fabrizio Mafessoni, Or Gross, Cathy Melamed-Bessudo, Shdema Filler-Hayut, Tal Dahan-Meir, Ziva Amsellem, Wojciech P. Pawlowski, Avraham A. Levy

## Abstract

The fate of DNA double-strand breaks (DSBs) generated by the Cas9 nuclease has been thoroughly studied. Repair via non-homologous end-joining (NHEJ) or homologous recombination (HR) is the common outcome. However, little is known about unrepaired DSBs and the type of damage they can trigger in plants. In this work, we designed a new assay that detects loss of heterozygosity (LOH) in somatic cells, enabling the study of a broad range of DSB-induced genomic events. The system relies on a mapped phenotypic marker which produces a light purple color (Betalain pigment) in all plant tissues. Plants with sectors lacking the Betalain marker upon DSB induction between the marker and the centromere were tested for LOH events. Using this assay we detected a flower with a twin yellow and dark purple sector, corresponding to a germinally transmitted somatic crossover event. We also identified instances of small deletions of genomic regions spanning the T-DNA and whole chromosome loss. In addition, we show that major chromosomal rearrangements including loss of large fragments, inversions, and translocations were clearly associated with the CRISPR-induced DSB. Detailed characterization of complex rearrangements by whole genome sequencing, molecular, and cytological analyses, supports a model in which breakage-fusion-bridge cycle followed by chromothripsis-like rearrangements had been induced. Our LOH assay provides a new tool for precise breeding via targeted crossover detection. It also uncovers CRISPR mediated chromothripsis-lke events that had not been previously identified in plants.

## Introduction

DNA double-stranded breaks (DSBs), can be repaired by non-homologous end joining (NHEJ), or by homologous recombination (HR). Error-prone, NHEJ can generate small insertions or deletions (Indels) at the DSB site (Gorbunova and Levy, 1997). The outcomes of HR vary according to the homologous partners: recombination between repeats in cis can lead to deletions in the case of direct repeats or inversions in the case of inverted repeats (Lupski, 1998). HR between ectopic repeats (Shalev and Levy, 1997; Puchta, 1999) can lead to translocations. DSBs were also shown to induce crossovers between sister chromatids or homologous chromosomes in somatic cells (Molinier et al., 2004). The CRISPR-Cas9 system enables analyzing the DSB repair process at endogenous loci and in a targeted manner, becoming an invaluable tool for precise breeding (Barrangou and Doudna, 2016). It is now possible to perform targeted mutagenesis, or when multiple breaks are induced, NHEJ enables precise chromosome engineering through deletions, inversions or translocations of large chromosomal segments (Beying et al., 2020; Schmidt et al., 2020). CRISPR-induced HR-mediated repair enabled enhancing gene replacement frequencies (Schiml et al., 2014; Baltes et al., 2014; Dahan-Meir et al., 2018) or achieving targeted crossovers or gene conversions (Filler-Hayut et al., 2017; Ben Shlush et al., 2020; Filler-Hayut et al., 2021). However, in the absence of selection, rates of targeted crossover are quite low (Filler-Hayut et al., 2021), and in tomato, targeted crossover events were so far identified only by using visual fruit color markers (Filler-Hayut et al., 2017; Ben Shlush et al., 2020).

While the promises of genome editing for precise plant breeding are immense, there are still many challenges. Repair is not always efficient and unrepaired DSBs can have deleterious consequences that have not been carefully analyzed in the CRISPR context. Recent studies in mammalian cells have demonstrated that CRISPR-Cas9 can induce loss of heterozygosity (LOH) as a result of loss of segments, arms, or whole chromosomes, as well as a cascade of chromosomal rearrangements following cell divisions (Zuccaro et al., 2020; Alanis-Lobato et al., 2021; Leibowitz et al., 2021). These rearrangements are similar to those found in cancer, and can have unintended consequences on the edited genomes and cells. A few percent of the cells that underwent Cas9-induced DSBs, exhibit chromosome bridges and, later, micronuclei (Leibowitz et al., 2021). This is consistent with a model in which chromosomal rearrangements occur through breakage-fusion-bridge-cycles (BFBC). Such cycles were first discovered in the pioneering work of Barbara McClintock, who studied the fate of chromosomes broken through irradiation or transposable element activities in maize. She proposed that sister chromatids of chromosomes with unrepaired broken ends become joined *via* what we call now NHEJ, generating a dicentric chromosome that can be broken at anaphase when the two centromeres are pulled to the opposite poles (McClintock, 1941). This process can repeat itself, triggering a cycle of random breaks between the centromeres and subsequent repair, leading to a breakage-fusion-bridge-cycle (BFBC). The outcomes of this process can be a range of chromosomal rearrangements, including deletions, duplication and inversion of large chromosomal segments or whole-chromosome loss. In cancer cells, the BFBC was shown to further trigger a series of catastrophic events, leading to chromothripsis: for example, an acentric segment can be excluded from the nucleus, forming a micronucleus, then the micronucleus DNA content can undergo fragmentation and the resulting fragments can re-integrate into the genome, causing additional chromosome rearrangements (Kwon et al., 2020; Ostapińska, Styka and Lejman, 2022).

In plants, there has been no evidence so far for the induction of a BFBC by CRISPR-Cas9-induced DSBs. In addition, chromothripsis has received little attention so far, except for findings in Arabidopsis, suggesting that genome elimination occurring in hybrids containing an altered centromeric histone CENH3 was reminiscent of chromothripsis (Tan et al., 2015; Henry, Comai and Tan, 2018).

In this work, we developed a new assay that enables visual detection of LOH using a hemizygous transgenic Betalain marker. DSB-induced LOH, due to somatic loss or homologous recombination of chromosomal segments carrying the Betalain marker, could be seen as green sectors in a purple background or as twin sectors (green and dark purple) in leaves. Betalains are purple pigments found in flowers, leaves, roots, and fruits of plants of most families of the Caryophyllales (Strack, Vogt and Schliemann, 2003). Three enzymes are essential in the biosynthesis pathway of Betalains, and a cassette containing genes encoding them was inserted into several plant species, including *Solanum lycopersicum* (tomato) (Polturak et al., 2016). The genomic integration site of the T-DNA construct containing the three genes of the Betalain purple pigment biosynthetic pathway was identified, and the CRISPR-Cas9 system was used to induce DSBs between the marker and the centromere. Somatic sectors corresponding to putative deletion or HR events were segregated or regenerated into whole plants and analyzed at the molecular level, identifying a flower somatic sector on chromosome 3. These observations validate the concept that rare somatic crossover events can be visually detected at a desired locus, using a nearby transgenic marker. In addition, the LOH assay enabled us to detect unrepaired DSB events on chromosome 11, leading to the loss of the whole chromosome or chromosome segments carrying the T-DNA marker. We show that these events can be explained by the induction of a BFBC leading to chromothripsis.

## Results

### A general system for DSB-induced LOH detection using a Betalain marker and CRISPR-Cas9

In order to characterize DSB-induced LOH events, and understand their underlying cause, we have developed a system based on a known-location dominant visual genetic marker (Betalain) in a heterozygous background (Figure 1). A DSB induced in somatic cells anywhere between the marker and the centromere can be repaired by NHEJ, with no phenotypic consequences, or by HR, which, in case of a crossover can yield a twin sector, i.e. a wild type (WT) color transgene-free sector and a transgene-homozygous dark purple sector in the light purple hemizygote background. If the DSB is unrepaired, LOH can also be detected as loss of the phenotypic marker.

**Figure 1.**
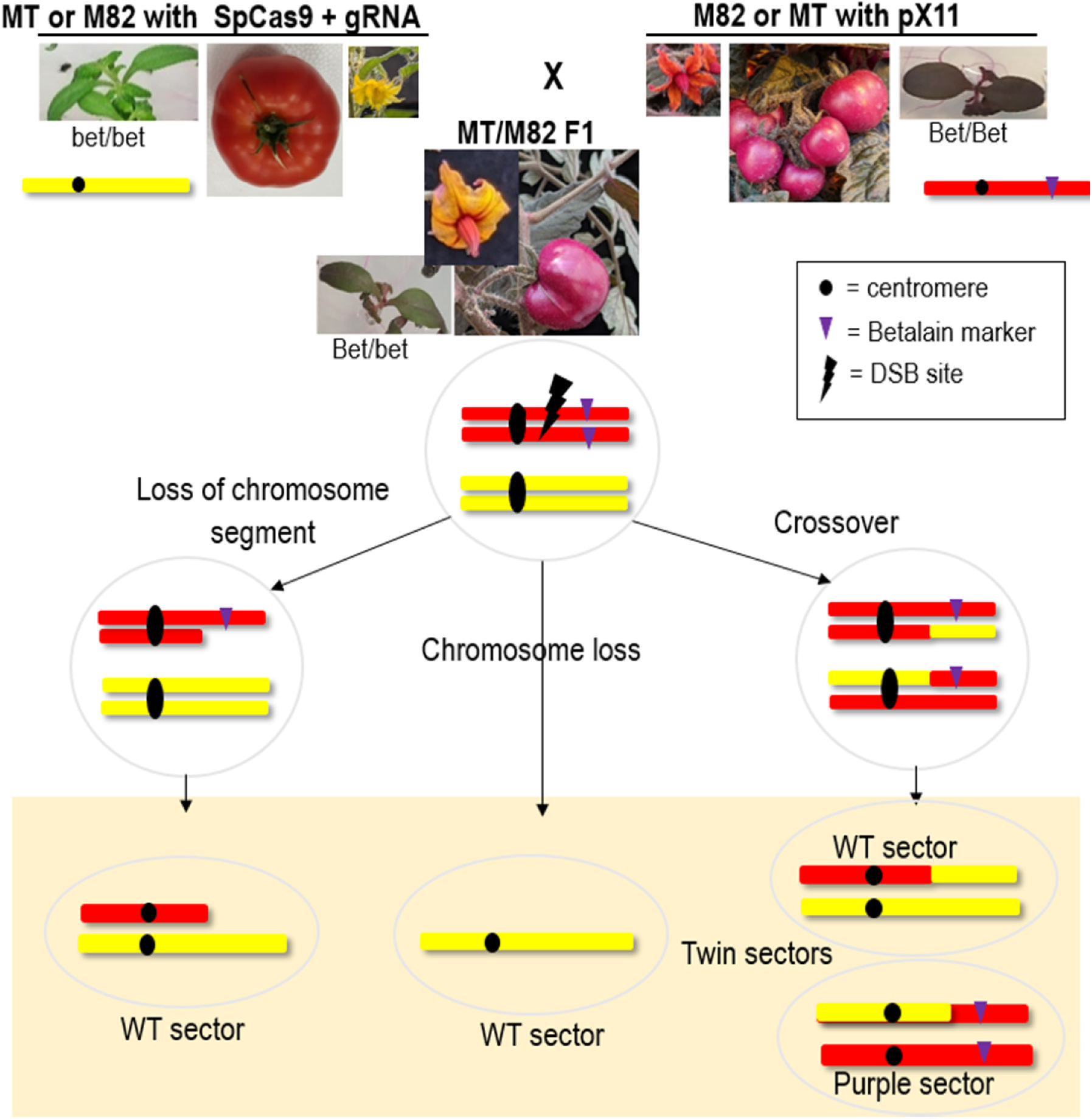
The Betalain visual assay for loss of heterozygosity (LOH) via deletion or crossover. Green MT/ M82 SpCas9 + gRNA is crossed with purple M82/ MT with Betalain (Bet) expression T-DNA (pX11) to generate an F1 hybrid. The light purple F1 is hemizygous for the pX11 T-DNA (Bet/bet). The chromosome on MT/ M82 SpCas9 + gRNA target line, has mutations at the gRNA target site and cannot be broken. The target chromosome in M82/ MT pX11 plants can undergo a DNA DSB at the target site. The DSB could be perfectly repaired or repaired with small indels by NHEJ. In these cases, the plants will remain light purple and will not be selected. LOH outcomes that can generate a WT phenotype (green leaf sector, yellow flower sector, or red fruit sector without Betalain) are shown in the bottom box. From left to right: loss of DSB distal fragment or part of it that contains the pX11 T-DNA, whole chromosome loss, or crossover that can generate a twin sector (WT color adjacent to dark purple). Black circles are representing the centromere. The purple triangle represents the T-DNA containing the Betalain marker and the black lightning bolt represents the site of CRISPR-Cas9 break.

The T-DNA construct (pX11) containing three Betalain biosynthesis genes was previously transformed into Micro-Tom (MT) (Polturak et al., 2017). Seeds were kindly provided by the Aharoni lab. We also generated new lines carrying pX11 in the M82 background. We mapped the T-DNA insertion site using inverse PCR (Figure 2). Primers were designed for the left border (LB) region of the pX11 cassette (Supplementary Table S1). DNA from pX11 homozygous lines in the M82 and MT backgrounds was extracted and then digested using PstI and HindIII restriction enzymes respectively (Figure 2A). The digested DNA was then self-ligated and subjected to two rounds of PCR amplification with nested primers. PCR products were Sanger-sequenced using the LB primer (Figure 2B) (Thomas et al., 1994). Sequence regions that did not align with the pX11 sequence were then BLASTed against the *S. lycopersicum* genome, which revealed the location of the junctions between the pX11 T-DNA and the M82 or MT genomes. Primers were then designed from both sides of the T-DNA insertion site to verify both the LB and RB junctions. This procedure confirmed the pX11 integration sites. In M82, the pX11 integration site was found on the short arm of chromosome 3, SL4.0ch3: 2,475,240, downstream of Solyc03g007960 (Figure 2C). In MT, pX11 integration was on the long arm of chromosome 11, SL4.0ch11:47305369, upstream of Solyc11g062370 (Figure 2D).

**Figure 2.**
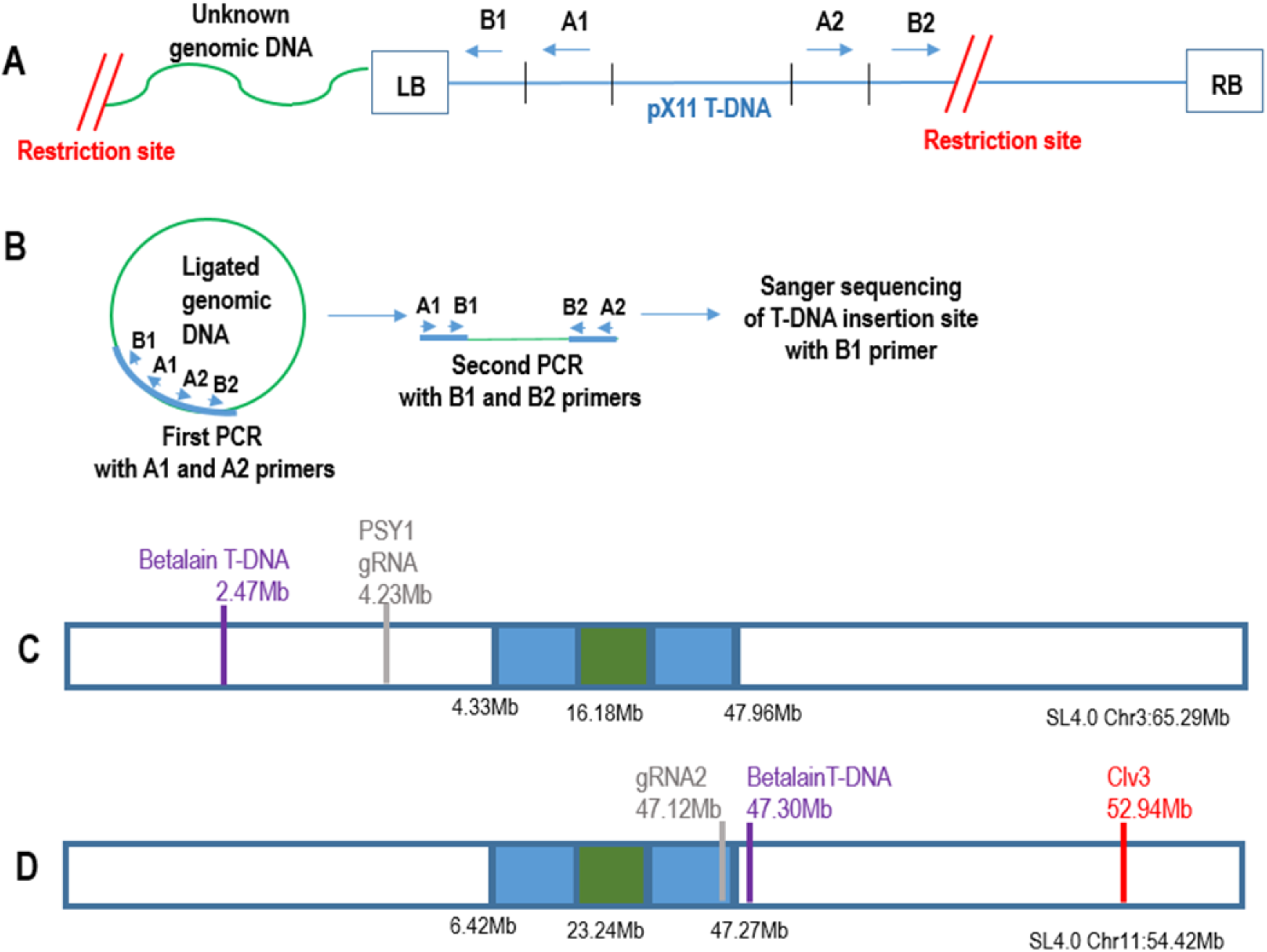
Sequencing pX11 T-DNA insertion sites by inverse PCR. A: Plant DNA is restricted with restriction enzyme (PstI or HindIII), followed by self-ligation. B: PCR amplification with two sets of nested primers, A1, A2 then B1, B2 followed by Sanger sequencing. This enabled the identification of the genomic sequence at the junctions of the Left border (LB) of the T-DNA. C: Illustration of SL4.0 chromosome 3 with coordinates of the PSY1 gRNA DSB (light gray line) target, and the mapped pX11 T-DNA integration site in M82 (purple line). D: Illustration of SL4.0 chromosome 11 with coordinates of the gRNA2 DSB target (light gray line), the mapped pX11 T-DNA integration site in MT (purple line), and the *CLV3* locus (red line). Centromere – green box; heterochromatic region – blue box; euchromatic region – white box. Estimated SL4.0 coordinates of centromere, heterochromatic and euchromatic regions are given below in mega bases (Mb). The complete size in Mb for each chromosome is shown underneath the chromosome illustration.

In order to identify LOH events, the system included the mapped Betalain marker (Figure 2) in the F1 hybrid (MT x cv M82-pX11) and (cv M82 x MT-pX11) backgrounds. LOH could be identified as dark purple Bet/Bet, or WT transgene free (green in leaves, yellow in flowers, or red in fruit) somatic sectors in the hemizygote light purple Bet/bet background (Figure 1). Moreover, SNPs differentiating between the parental lines could be used for genotyping at the whole-chromosome scale. Targeted DSBs were induced with CRISPR-Cas9 between the centromere and the T-DNA. MT and M82 plants containing the SpCas9 + gRNA, both controlled by constitutive promoters, were crossed to M82 and MT pX11 plants, respectively (Figure 1).

On chromosome 3, we designed the gRNA to target a euchromatic region (Demirci et al., 2017) at position SL4.0ch3:4,234,645 in exon4 of *PSY1*, at the distance of 1,759,405 bp from the M82 pX11 marker towards the centromere (Figure 2C, Supplementary Table S2). A transgenic MT plant carrying SpCas9 and *PSY1* gRNA produced an NHEJ footprint at the DSB target of GCT deletion (-GCT) in 50% of the reads, and G deletion (-G) in 50% of the reads (Dahan Meir et al., 2018). This represents one chromosome with (-GCT) and one chromosome with (-G) footprints. Progeny of this MT plant carrying SpCas9 and *PSY1* gRNA, produced by selfing had the same NHEJ footprints at the *PSY1* gRNA DSB target site, [100% (-GCT), or 100% (-G), or 50% (-GCT)/ 50% (-G)], indicating no further DSB formation. The transgenic MT plant carrying SpCas9 and *PSY1* gRNA was used for crossing with M82 pX11. Since the *PSY1* gRNA target on the MT chromosome 3 was mutated, only the M82 pX11 chromosome 3 could be cleaved by Cas9 in the F1 plants.

On chromosome 11, we targeted the gRNA to a heterochromatic region (Demirci et al., 2017) at position SL4.0ch11: 47,124,456 (gRNA2), between two gene promoters at the distance of 180,895 bp from the MT pX11 marker towards the centromere (Figure 2D, Supplementary Table S2). The transgenic M82 plant with SpCas9 and gRNA2 gave an NHEJ footprint at the DSB target of +T insertion (+T) in 100% of the reads. Progeny of this plant produced by selfing had the same NHEJ footprint at the chromosome 11 gRNA2 target site [100% (+T)], indicating no further DSB formation. The transgenic M82 plant with SpCas9 and gRNA2 was used for crossing with MT pX11. Since the gRNA2 target of the M82 chromosome 11 was mutated, only the MT pX11 chromosome 11 could be cleaved by Cas9 in the F1 plants.

### Screening for twin sectors as putative crossover events

Twin sectors, consisting of a dark purple sector (*Bet/Bet*) adjacent to a WT sector (*bet/bet*) in the light purple background (*Bet/bet*) can represent putative somatic crossover events originating from a reciprocal exchange between chromatids of homologous chromosomes as shown in Figure 1. To induce and visually identify such somatic events, we screened (*Bet/bet*) plants where DSBs were induced between the Betalain markers mapped on chromosomes 3 and 11 and centromeres as shown in Figure 2 C and D, respectively.

To search for twin sectors for the chromosome 3 target (Figure 2C), ten F1 light purple (*Bet/bet)* plants containing SpCas9 + PSY1 gRNA and ten F1 purple (*Bet/bet*) control plants containing SpCas9 but lacking the gRNA (Supplementary Table S3) were grown in the greenhouse. Most plants appeared to be phenotypically heterozygous (light purple) with no twin sectors large enough to represent a high-confidence CO event (Figure 3A). A flower of one of the plants showed in a petal a somatic twin sector consisting of a dark purple *Bet/Bet* sector adjacent to a yellow bet/bet (WT) sector in the background of light purple *Bet/bet* (Figure 3B, Supplementary Table S4). The fruit generated from this chimeric flower produced six viable F2 progeny plants. Among them, one light purple F2 (*Bet/bet*) plant had a crossover (CO) event on chromosome 3. On the telomere side, both the *PSY1* gRNA site and the SNP closest to the *PSY1* gRNA were heterozygous (M82/MT) while the next SNP towards the centromere was homozygous MT (Figure 3C, 3D). This plant was analyzed by whole genome sequencing and did not exhibit changes in the number of reads along both sides of the DSB site (Supplementary Figure S1). Therefore, this change from heterozygote to homozygote is unlikely to be due to a chromosome segment loss. The plant was fertile and produced viable F3 seeds further confirming that this was a genuine targeted CO. We did not find CO events in ten F2 progeny of other fruits of the same plant, nor in ten F2 progeny of M82 pX11 x MT SpCas9 control plants lacking *PSY1* gRNA.

**Figure 3.**
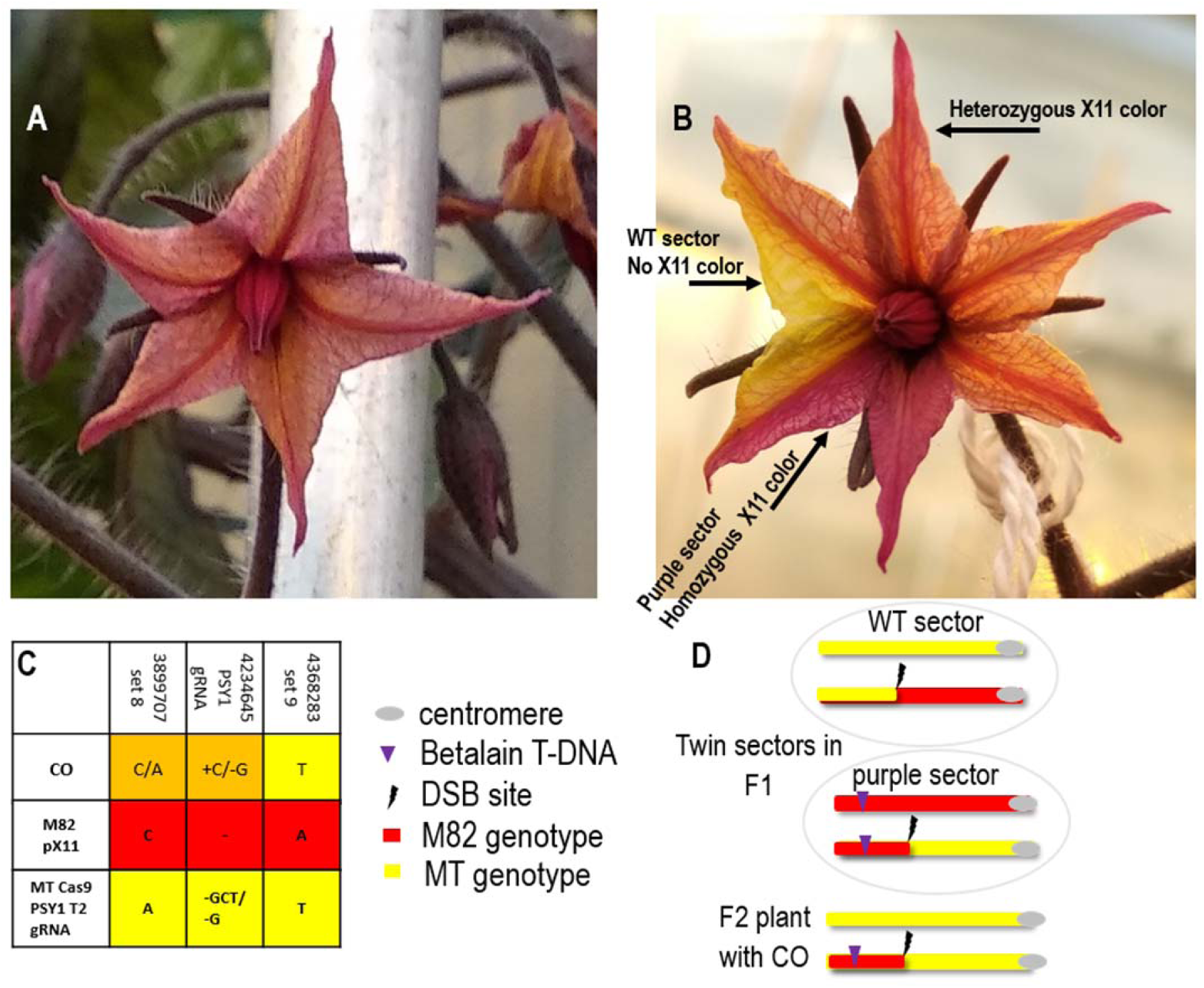
Detection of a targeted somatic crossover event on chromosome 3 seen as a twin sector. A DSB was induced by the *PSY1* gRNA on chromosome 3, between the centromere and the Betalain marker. A: Tomato flower of F1 plant [(MT SpCas9 + PSY1 gRNA)/ (M82 pX11 on chromosome 3)] with uniform light purple color phenotype. B: Tomato flower of F1 plant [(MT SpCas9 + PSY1 gRNA)/ (M82 pX11 on chromosome 3)] with chimeric twin-sector color phenotypes (WT yellow (no transgene) next to a dark purple sector (Bet/Bet). One F2 progeny out of six in the fruit generated from this flower was a CO event. C: Sanger sequencing results of three SNPs sets on chromosome 3: 1) Set 8 upstream to the DSB site towards the telomere side; 2) SpCas9 + PSY1 gRNA, generated SNP at the DSB site in the MT SpCas9 + PSY1 gRNA background; 3) Set 9 downstream to the DSB site towards the centromere side. In the CO event F2 generated from the twin sector flower in the B panel, we see a transition from heterozygous MT/M82 SNPs in set 8 and PSY1 gRNA to homozygous MT SNPs in set 9. Orange highlight – heterozygous MT/M82 SNPs; red highlight – homozygous M82 SNPs; yellow highlight homozygous MT SNPs. D: Scheme of the recombinant chromosome 3 generated by SpCas9 induced somatic crossover, and detected by following the Betalain color marker. The F1 somatic recombination event generated twin sectors as seen in panel B. The putative chromosome 3 genotypes of each sector are presented in panel D. In the viable F2 plant with CO event, a gamete containing the chromosome 3 somatic CO product of the purple sector paired with a gamete containing the chromosome 3 MT parental type. Grey dot – centromere. Purple arrow Betalain T-DNA integration site. Black lightning bolt – DSB site. Orange highlight – heterozygous M82/MT SNPs; red highlight – homozygous M82 SNPs; yellow highlight homozygous MT SNPs.

For Chromosome 11 (Figure 2D), where the pX11 cassette was mapped to SL4.0ch11:47305369, we found an additional visual marker on the long arm of chromosome 11: the *CLAVATA3* (*CLV3)* gene located at position SL4.0ch11:52945095, between the pX11 cassette and the telomere. Seeds of the *clv3-1* mutant in the M82 background were kindly provided by Zach Lippman’s lab. The *clv3* mutants exhibit a fasciated phenotype in flowers and fruits (Xu et al., 2015), seen as an increase in floral organs and fruit locules compared to WT plants (Figure 5 M82 *clv3*). We used *clv3* as an additional genetic marker to detect recombination or other chromosomal rearrangements. *CLV3* is 5.82Mb upstream from gRNA2 towards the telomere. The transgenic M82 *clv3* plant with SpCas9 and gRNA2 was crossed with MT pX11. Ten F1 light purple (*Bet/bet; Clv3/clv3*) plants containing SpCas9 + gRNA2 and ten F1 purple (*Bet/bet; Clv3/clv3*) control plants containing SpCas9 but lacking the gRNA (Supplementary Table S3), were grown in the greenhouse. After 8 weeks, all plants appeared phenotypically heterozygous to the Betalain marker (light purple) with no visible twin sectors large enough to represent COs and no green sectors expected of LOH events.

### Genotyping and phenotyping of Somatic LOH events

As the presence of large sectors is a rare occurrence, we wanted to have an additional screen that allows the detection and regeneration of whole plants from small sectors. To do it, we grew ten F1 seedlings each for chromosome 3 and 11 markers in sterile conditions for regeneration from cotyledons to search for small green somatic sectors. Our plan was to regenerate whole plants from these sectors and identify the underlying cause of their phenotypes. We hypothesized that the F1 plants either experienced loss of heterozygosity (LOH) or that the transgene was silenced. Two-week-old purple seedlings (Bet/bet), confirmed for SpCas9 presence (Supplementary Table S3), were dissected into small pieces and transferred into tissue culture for whole-plant regeneration. On average 160-180 leaf pieces per plant were prepared from each of the ten F1 purple (Bet/bet) plants. Calli and newly emerging plantlets were identified, and green plantlets were observed (Figure 4).

**Figure 4.**
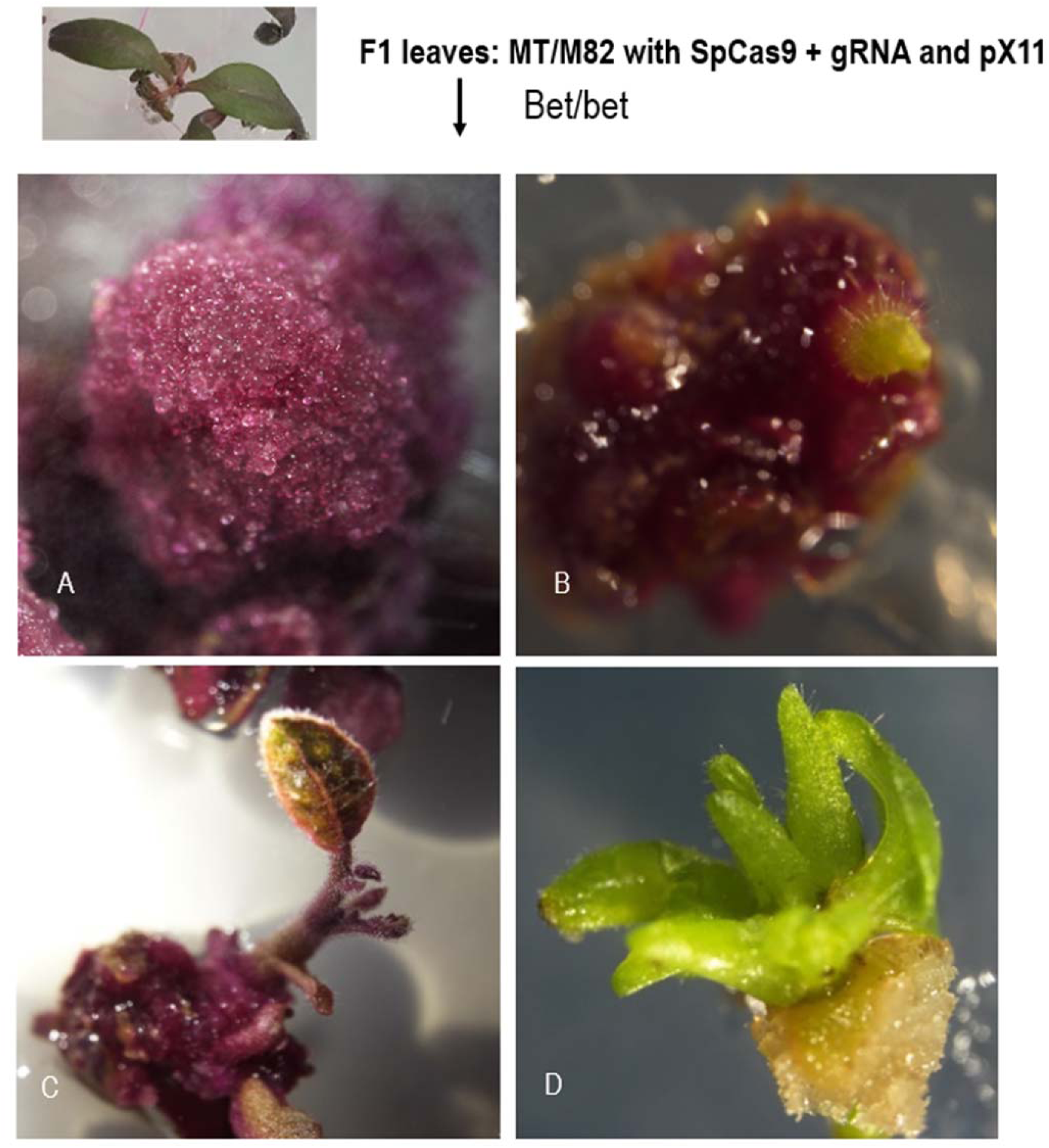
Screening for green sectors regenerated from F1 somatic tissues. We analyzed F1 light purple and SpCas9 positive plants of the genotypes [(MT SpCas9 + PSY1 gRNA)/ (M82 pX11 on chromosome 3)], or [(M82 *clv3* SpCas9 + gRNA2)/ (MT pX11 on chromosome 11)]. Green and light purple plantlets regenerated from pieces of light purple F1 leaves were obtained. A light purple callus (A) and a light purple plantlet (C) were regenerated from a light purple F1 leaf. Green calli (B) occasionally emerged from a purple callus giving rise to a green regenerated plantlet (D). Green plantlets were further analyzed as potential LOH products.

**Figure 5.**
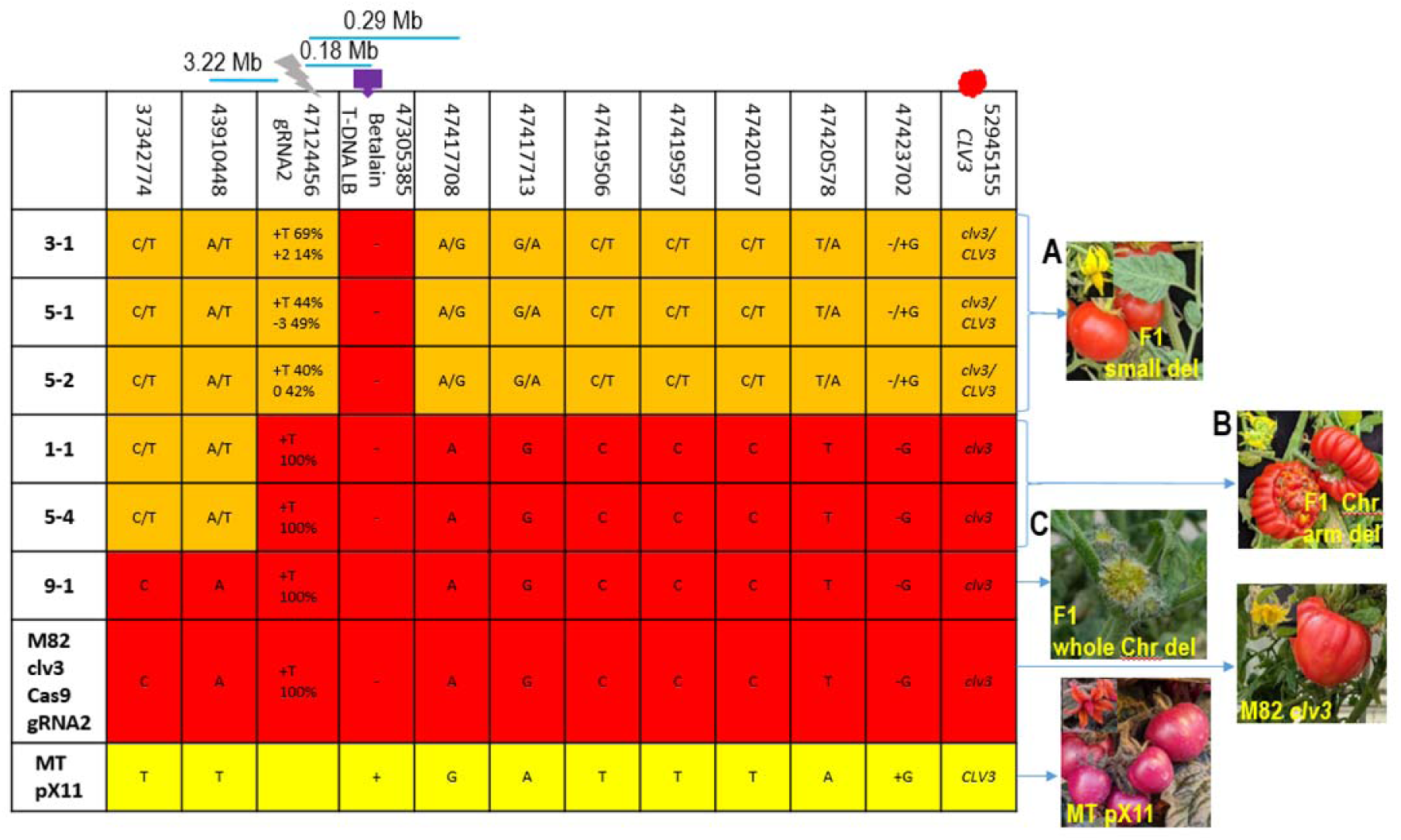
SNP genotyping of regenerated F1 green plant. Six regenerated green plantlets derived from F1 hybrids and their parents were sequenced for SNP markers in the region around the chromosome 11 gRNA2 induced DSB and for the *CLV3* marker genotype and phenotype. Coordinates of SNPs on SL4.0 chromosome 11 are noted in the top row of the table. The DSB point is marked as a grey lightning bolt. A: In plants 3-1, 5-1, and 5-2 the T-DNA was missing. SNPs were heterozygous for the genotype of both parents, upstream and downstream of the DSBs all the way to the *CLV3* gene, which is located ∼2 Mb upstream from the telomere, as expected for an F1. Flowers and fruit had a *CLV3* WT phenotype. B: In plants 1-1, and 5-4, the T-DNA was missing. Upstream of the DSB point, SNPs were heterozygous. After the DSB and all the way to the *CLV3* gene, SNPs were homozygous M82. Flowers and fruit had a *clv3* fasciated phenotype. C: In plant 9-1, The T-DNA was missing. Upstream and downstream of the DSB, SNPs, including the *CLV3* gene, were homozygous for M82. Flowers had an extreme *clv3* fasciated phenotype, and no fruits were generated. Orange highlight – heterozygous M82/MT SNPs; red highlight – homozygous M82 SNPs; yellow highlight homozygous MT SNPs. Chr-chromosome; del – deletion. 3.22Mb – distance between gRNA2 and the first SNP downstream towards the centromere. 0.18Mb – distance between the Betalain T-DNA integration site and gRNA2. 0.29Mb – distance between the first SNP upstream towards the telomere and gRNA2. Grey lightning bolt – DSB site. Purple arrow Betalain T-DNA integration site. Red dot – *CLV*3 position.

For each green regenerated plant, the presence/absence of the pX11 T-DNA, or at least one of its borders junctions with the genomic integration site, was verified by PCR. For chromosome 3, two F1 green plants were regenerated, representing candidates for LOH. We found that the plants did not lose the pX11 T-DNA, thus we concluded they were silencing events. One silencing event was also observed among the control F1 purple (*Bet/bet*) SpCas9 plants. For chromosome 11, one silencing event was observed among F1 purple (*Bet/bet*; *CLV3/clv3*) SpCas9 + gRNA2 plants and one in the control among F1 purple (Bet/bet; *CLV3/clv3*) SpCas9 plants. These data suggest that gene silencing was not related to the occurrence of distant DSBs. Thus, these silencing events (Supplementary Table S4) were not further analyzed.

In addition to the plants carrying gene silencing events, other green regenerated plants were analyzed (Figure 5, Supplementary Table S4). Three such plants (Figure 5; plants 3-1, 5-1, 5-2) had lost the T-DNA insertion region but were heterozygous for all SNPs around the T-DNA, including in the region of the gRNA target, as expected for F1 plants (Figure 5). Moreover, these plants also had the expected F1 *Clavata* genotype (*Clv3/clv3*) and phenotype (Figure 5A) and were fertile, suggesting that no major chromosomal rearrangements had occurred.

Two plants lacked the T-DNA pX11 cassette and showed a transition from heterozygous SNPs to SNPs homozygous for the M82 parent (Figure 5, plants 1-1, 5-4) in the DSB region. The plants were also homozygous for the telomeric *clv3* allele mutation and had severely fasciated flowers and fruits (Figure 5B). These plants could have been considered exhibiting targeted crossover events, based on their SNP genotype and the phenotype. However, these phenotypes and genotypes could also be explained by a loss of chromosome arm from the DSB site to the telomere. This possibility was strengthened by the high sterility of the plants and was further confirmed through whole genome sequencing (see below) (Figure 6).

**Figure 6.**
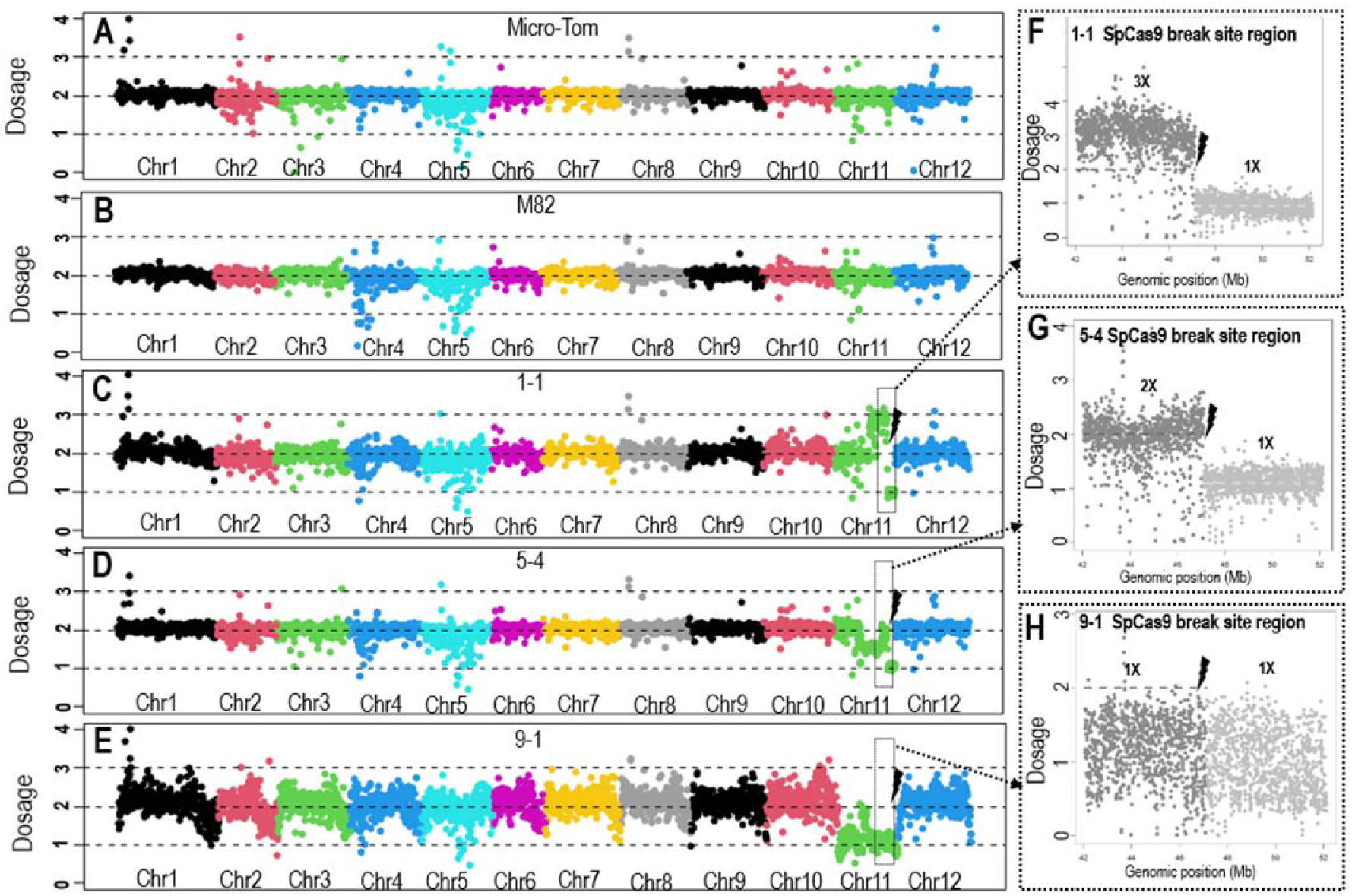
Dosage changes at the induced DSB site by WGS coverage analyses. Average coverage of WGS reads per plant was determined as 2X diploid dosage basis. A-E: Coverage for each of the 12 chromosomes per plant, is presented with each chromosome shown in a different color. Panel A and B: parental plants with 2X in all chromosomes. Panel C: plant 1-1 showing deviation of 2X ploidy in chromosome 11. Panel F: Plant 1-1 zoom-in on chromosome 11, about 5Mb from each side of the gRNA2 DSB site. The dosage in each region reveals changes in ploidy levels. Plant 1-1 shows transition at the DSB site, from a dosage of 3X to 1X. This indicates loss of the region from the DSB site to the telomere in one of the chromosomes. Panel D: plant 5-4 showing deviation of 2X ploidy in chromosome 11. Panel G: Plant 5-4 zoom-in on chromosome 11, about 5Mb from each side of the gRNA2 DSB site. The dosage in each region reveals changes in ploidy levels. Plant 5-4 shows transition at the DSB site, from a dosage of 2X to 1X. This indicates loss of the region from the DSB site to the telomere in one of the chromosomes. Panel E: Plant 9-1 shows a dosage of 2X along all chromosomes but throughout chromosome 11 there is a dosage of 1X indicating loss of the whole chromosome from one of the parents. Panel H: Plant 9-1 zoom-in on chromosome 11, about 5Mb from each side of the gRNA2 DSB site. The dosage in each region reveals no change in the ploidy levels showing a dosage of 1X on both sides of the DSB site. Chr-chromosome. Black lightning bolt – DSB site.

Plant 9-1 also lacked the T-DNA pX11 cassette and was homozygous for all the M82 SNPs. Moreover, it was homozygous for the *clv3* allele it had the most severely fasciated flower phenotype and was totally sterile, bearing no fruits (Figure 5C). This genotype/phenotype could be explained by a complete chromosomal loss, as confirmed by whole genome sequencing analysis described below (Figure 6).

### Whole genome sequence analysis of LOH events

Whole genome sequencing (WGS) was performed for plants 3-1, 5-1, 5-2, 1-1, 5-4, 9-1, and for the Micro-Tom and M82 lines. The sequencing coverage, in plants 3-1, 5-1, and 5-2 (Supplementary Figure S2A, S2B, S2C, respectively), was similar to that of the parental plants, Micro-Tom and M82 (Figure 6A, 6B). The ploidy dosage was determined using sequencing coverage analysis as described in the Materials and Methods. SNPs in the reads could be anchored to the parental sequences and therefore this analysis enabled assessing deviations from the diploid dosage as well as homozygosity, heterozygosity, and hemizygosity.

Analysis of F2 progeny of plant 5-1 homozygous for MT SNPs on both sides of gRNA2 and pX11 revealed a 4069bp fragment between the NOS terminator inverted repeats was missing from the pX11 T-DNA on chromosome 11 (Supplementary Figure S3A). Except for the deleted part of the pX11 T-DNA in plants 3-1, the dosage was 2X in all chromosomes, including chromosome 11.

WGS of F2 progeny of plants 3-1 and 5-2, which were also homozygous for MT SNPs on both sides of gRNA2 and pX11, showed that a 17396bp fragment of chromosome 11 (SL4.0ch11:47301145 -47318550) spanning the original T-DNA insertion site (SL4.0: 47305385) was missing, as confirmed by Sanger sequencing (Figure 5). Except for the deleted parts spanning the pX11 T-DNA in plants 5-1, and 5-2 the dosage was 2X in all chromosomes, including chromosome 11. Both sides of the deleted region spanning the T-DNA, are flanked by A-rich repeats (Supplementary Figure S3B). Note that T-DNA loss was not detected in control plants where no DSB induction occurred.

WGS of F1 9-1 showed the expected 2X coverage, and SNP heterozygosity, throughout the genome except for chromosome 11 where significant deviations were observed (Figure 6E). The coverage dosage was 1X throughout chromosome 11, and SNPs corresponded only to the M82 parent (Figure 6E). Note that such massive loss was detected only in plants containing Cas9 and gRNA where a DSB was induced, and not in control plants.

WGS of plants 1-1 and 5-4 confirmed the transition from heterozygous to M82 SNPs precisely at the SpCas9-induced DSB site (Figure 5). The analysis of dosage of sequencing reads showed that in both 1-1 and 5-4, the dosage around the targeted DSB site changed from 3X (possible duplication) or 2X (diploid), respectively, to 1X (haploid). This result indicates that a segment from the DSB region closer to the telomere of the long arm of chromosome 11 was lost (Figure 6C, 6D, 6F, 6G) as it would be expected for unrepaired DSB events.

WGS of F1 9-1 showed the expected 2X coverage, and SNP heterozygosity, throughout the genome except for chromosome 11 where significant deviations were observed (Figure 6E). The coverage dosage was 1X throughout chromosome 11, and SNPs corresponded only to the M82 parent (Figure 6E, 6H), with the exception of reads across the centromere, possibly due to mapping biases. Note that such massive loss was detected only in plants containing Cas9 and gRNA where a DSB was induced, and not in control plants.

### Multiple chromosomal rearrangements are associated with telomere loss

The WGS dosage analysis revealed additional rearrangements in plants 1-1 and 5-4. Plant 1-1 had a ∼20Mb region between the DSB and the centromere that became duplicated, showing a 3X chromosome dosage (Figure 7). This duplication transitioned to chromosome segment showing a 1X dosage precisely at the DSB site (Figure 7A). This kind of duplication and deletion event could be explained by loss of an acentric chromosome fragment distal to the DSB (and hence the 1X dosage) followed by fusion of two centromere-bearing sister chromatids, which would initiate a breakage-fusion-bridge-cycle (BFBC) (Figure 7D). During a BFBC, when the two centromeres are pulled to the opposite poles in anaphase, a new break is generated randomly along the bridge, eventually leading to a duplication, hence the 3X dosage shown as region A (Figure 7A and D) in an inverted orientation (Figure 7D).

**Figure 7.**
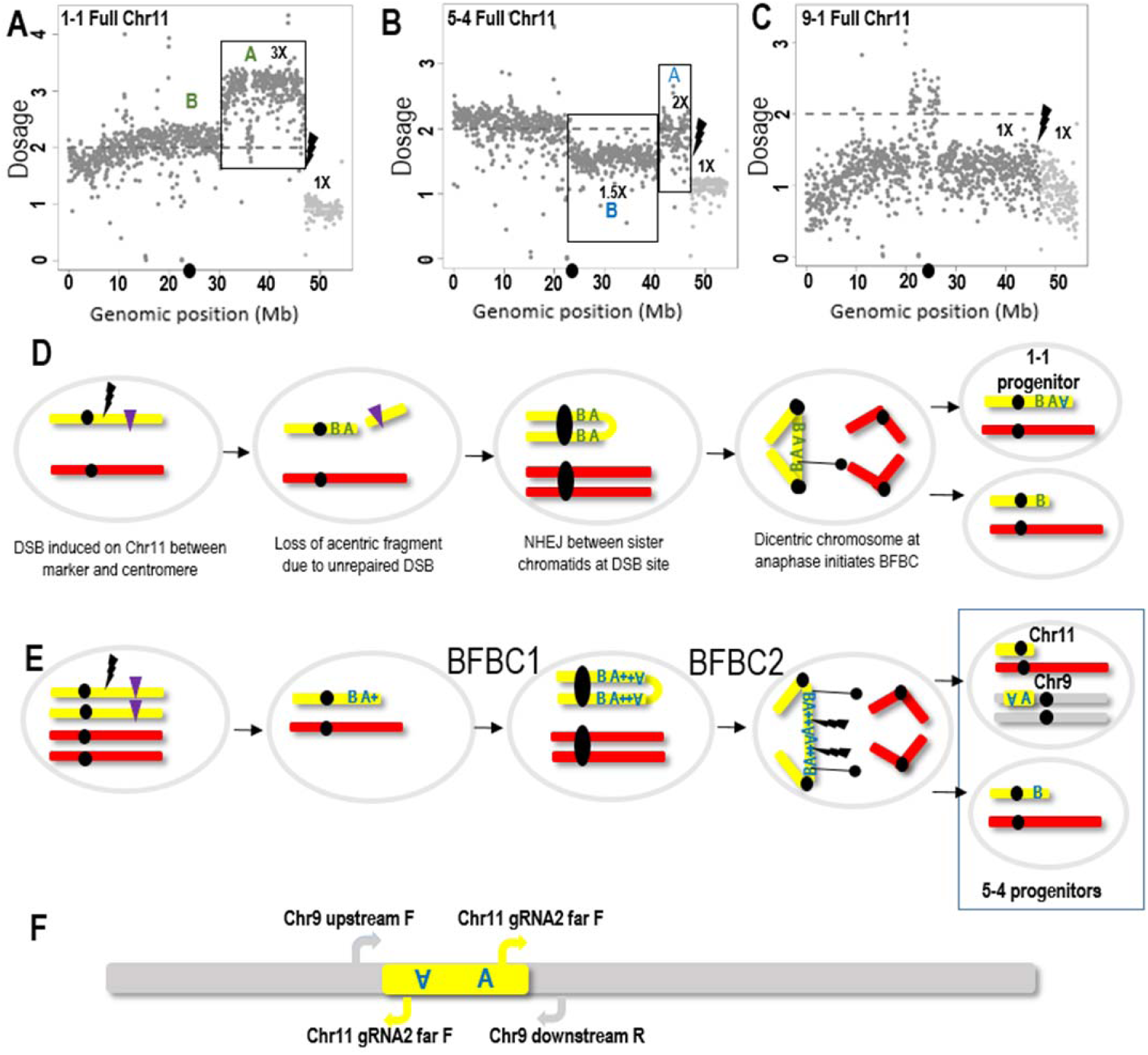
Analysis of chromosomal rearrangements at DSB sites and an underlying mechanistic model of breakage fusion bridge cycle (BFBC) and chromotripsis. A: Plant 1-1 whole chromosome 11 coverage shows transition at the DSB site, from a dosage of 3X to 1X. This indicates loss of the region from the DSB site to the telomere in one of the chromosomes and duplication in the other side of the DSB marked by a grey A. B: Plant 5-4 whole chromosome 11 coverage reveals changes in ploidy levels, which are more complicated compared to plant 1-1. Plant 5-4 shows transition at the DSB site, from a dosage of 2X in the region marked by blue A to 1X from the DSB site to the telomere. This indicates loss of the region from the DSB site to the telomere in one of the chromosomes. An additional change from the average coverage is seen in the region marked by a blue B, which has a dosage of 1.5X. C: Plant 9-1 whole chromosome 11 coverage. In this plant, the dosage is 1X throughout the whole chromosome 11, except in the centromere region. D: Illustration of a putative BFBC that could explain the 3X dosage of the region marked by a grey A in plant 1-1. E: Illustration of putative two rounds of BFBC that could explain the 2X dosage of the region marked by a blue A and the 1.5X dosage of the region marked by blue B in the plant 5-4. The first round is similar to the panel D illustration. But here maybe the DSB occurred in the stage where two chromatids are present. The broken chromatid may have invaded the intact chromatid and copied the region containing the gRNA2 target site marked by +. In the second round, the bi-centric chromosome may have broken at several points. A duplicated part of the region marked by blue A + that contains the inverted duplicated A region and the gRNA2 target site from each side of the duplication, may have been cut by the SpCas9 and translocated into chromosome 9. F: Scheme of the chromosome 11 segment translocated into chromosome 9. We used two PCR primer sets to amplify and sequence the junctions. Chr9 upstream F + Chr11 gRNA2 far F for the upstream junction. The amplification of this junction using the Chr11 gRNA2 far F primer indicates the inverted orientation of the chromosome 11 blue A region. Chr11 gRNA2 far F and Chr9 downstream R used to amplify the downstream junction indicates the forward orientation of the chromosome 11 blue A region. MT chromosome 11 is highlighted in yellow; M82 chromosome 11 in red; Chromosome 9 in grey. Black lines with a black circle indicate a random DSB site generated by pulling of the bridge part to two opposite poles in the dicentric chromosome. The black lightning bolt indicates the SpCas9 DSB site. The chromosome 11 centromere is at 23.24 Mb, and is marked by a black dot on the Genomic position (Mb) axis.

In plant 5-4, a region between the DSB and the centromere was missing (Figure 7B). The estimated dosage in this region was 1.5X, which may indicate chimerism. To better understand this event, we examined the DSB junctions using inverse PCR amplifying sequences adjacent to the chromosome 11 gRNA2 region (Supplementary Table S6). This approach was successful in plant 5-4, but not in plant 1-1. Surprisingly, the amplified product showed a junction between the chromosome 11-gRNA2 and regions located on chromosome 9 (Figure 7F). We then designed a primer to the putative chromosome 9 sequence and verified the junction by PCR and Sanger sequencing (Supplementary Table S7, Supplementary Figure S4A). We also verified the upstream junction of the chromosome 11 gRNA region using a primer targeting the putative upstream chromosome 9 sequence and the primer from the chromosome 11-gRNA2 region (Figure 7F, Supplementary Figure S4B). The translocation represented by the two junctions was further confirmed by the finding of WGS Illumina reads corresponding to the chimeric junctions between chromosomes 11 and 9 (Supplementary Figure S5). The directionality of the joined Illumina reads, and the primers used for PCR amplification of the junctions support the model shown in Figure 7E, suggesting that in 5-4 the A inverted repeat region was broken and translocated to chromosome 9 (Figure 7E, 7F, Supplementary Figure S4, S5). Such rearrangement is similar to events resulting from chromothripsis (Ostapińska, Styka and Lejman, 2022; Stephens et al., 2011). We did not find any potential SpCas9-gRNA2 target sites in the chromosome 9 translocation region that could explain why the A fragment was inserted specifically at that location. Taken together, the complex rearrangements in plant 5-4 might be best explained through a scenario where the first BFBC round, which was similar to that in plant 1-1 (Figure 7d), was followed by a second round in which the A duplicated region was broken and translocated to chromosome 9. The second daughter cell would contain the broken MT chromosome 11 with the blue B region. These two daughter cells generated after the second BFBC may be the progenitors of the tissues analyzed in plant 5-4 (Figure 7E).

In plant 9-1, the coverage dosage was 1X throughout chromosome 11, with SNPs corresponding only to the M82 parent (Figure 7C), except for the centromere region (23.24Mb), where the dosage was 2X. Since the centromere region is rich in repeated sequences, it is difficult to determine if this is a misalignment artifact and in fact the whole MT chromosome was lost. Alternatively, it might be a near-complete loss of chromosome 11, with only a minichromosome consisting of chromosome 11 centromere remaining.

Cytological analysis of plants 1-1 and 5-4 pollen mother cells revealed the presence of bridges during meiotic anaphase I (Figure 8A), and micronuclei in tetrads (Figure 8B and 8C). The detection of bridges further suggests the presence of the chromosomal rearrangements detected by WGS. The rearrangements could be the results of prior BFBC episodes. The presence of micronuclei is also consistent with either the presence of acentric chromosome fragments or chromosomes that cannot pair properly, presumably as the result of their rearrangements.

**Figure 8.**
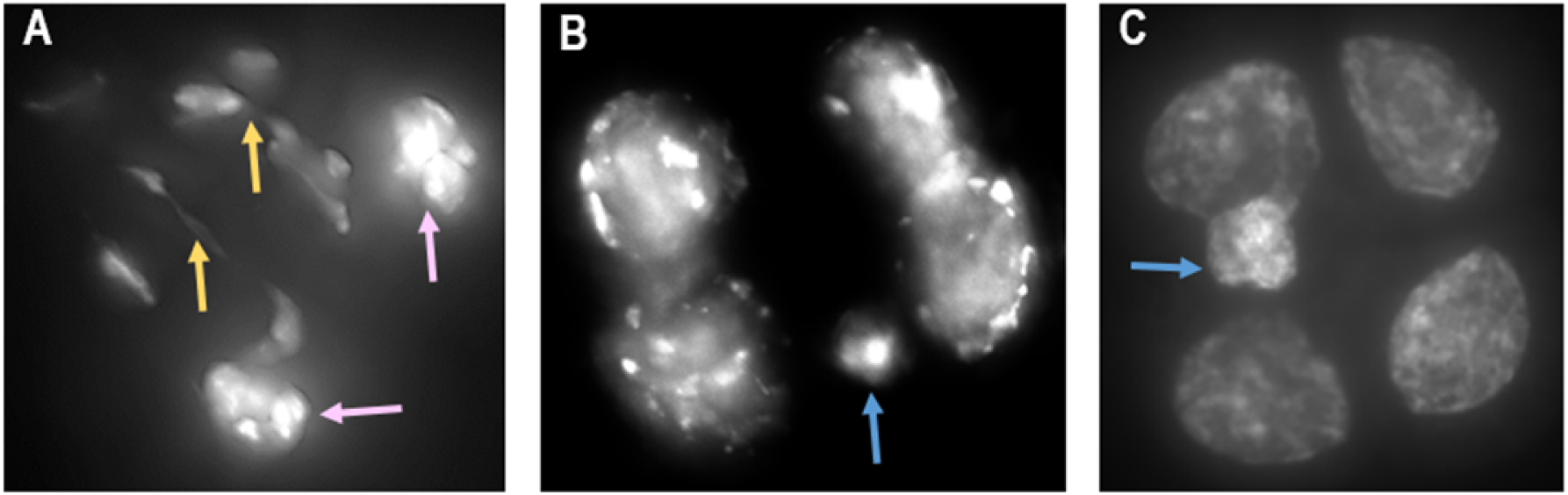
Cytological evidence of chromosomal rearrangements and micronuclei in meiocytes. A: Plant 1-1 meiocyte at late anaphase I showing two chromosome bridges (yellow arrows) and two clumps of chromosomes that have not properly segregated to daughter cells (pink arrows). B: Plant 1-1 tetrad with a micronucleus (blue arrow). C: Plant 5-4 tetrad with a micronucleus (blue arrow). Meiocytes from young flower buds were fixed in 3:1 ethanol: acetic acid and stained with DAPI.

## Discussion

We have developed a system that enables visual identification of a wide range of chromosome alterations. We report examples of such alterations, including a crossover and LOH due to chromosomal rearrangements ranging from segment to whole chromosome loss or translocation. The system is based on a previously-mapped visual transgenic marker. In principle, the Betalain biosynthesis marker can serve to monitor LOH on any chromosome as well as in many plant species, making this system quite general. This marker can be combined with additional phenotypic markers, as done here with the recessive *clv3* mutation on chromosome 11, to facilitate the analysis of the studied events.

We showed that twin sectors could be useful in detecting a targeted crossover at the site of induced DSBs located between the centromere and the marker on chromosome 3 (Figure 3). We found a single large twin sector in a flower of a plant out of 10 plants, and we confirmed in the progeny of this flower that a targeted crossover took place at the DSB induction site. Such crossover could give rise to reciprocal events, either a purple (i.e., containing the T-DNA marker) or a green WT transgene-free targeted recombinant. In this single event, we obtained a purple plant, but in principle, the system can be used also for the recovery of transgene-free germinal recombinant events making it a useful tool. The potential benefits of such systems in plant breeding have been thoroughly discussed (Taagen et al., 2020; Sukegawa, Saika and Toki, Rönspies et al., 2021). The rate of occurrence of twin sectors was low, which is consistent with our previous works on targeted somatic crossovers in tomato (Filler-Hayut et al., 2017; Ben Shlush et al., 2020) and Arabidopsis (Filler-Hayut et al., 2021). The system described here is not restricted to a particular chromosomal site and can be applied to screen for rare events. In addition, it can be used to screen for regions in the genome, or for mutants, that are more prone to repair via HR and via CO. Before, we used a natural marker at the *Sulfur* locus in tobacco to identify the *Hyrec* mutant exhibiting high somatic recombination rates (Gorbunova et al., 2000). The *Hyrec* locus has not been characterized at the molecular level, but its phenotype shows that reaching high rates of somatic crossover might be feasible.

Because of the low number of large green or twin sectors seen in whole plants, and to enable obtaining of germinal from somatic events, we also screened for green calli regenerated from leaves of light purple F1 plants. When testing the chromosome 3 euchromatic target, we did not detect WT sectors corresponding to DSB-induced LOH. With the heterochromatic chromosome 11 DSB target, we found 6 independent LOH events. All of these events were deletion events of various sizes and not CO events. The abundance of repeats in the chromosome 11 DSB target might have triggered defective repair as previously reported for centromeric regions (Barra and Fachinetti, 2018). These LOH events were genotyped locally around the break and genome-wide, using WGS. In three out of these 6 plants, the Betalain-coding T-DNA was fully or partially eliminated. The mechanistic underpinnings of these transgene elimination, 180 Kb away from the DSB, events are not clear. It is well known that somaclonal variation induced in tissue culture can give rise to chromosomal rearrangements, even without DSB induction. However, we did not detect T-DNA loss in SpCas9 control plants. It is possible that the deletions were induced through a long-range repair mechanism that remains to be discovered. Note that off-site DSBs are an unlikely explanation for T-DNA loss as deletions occurred in regions with no homology to the target, namely the NOS inverted repeats and an A-rich region.

Regarding the chromosome 11 loss in plant 9-1, it is conceivable that a DSB induced BFBC could lead to a very small and unstable chromosome, and this is further supported by the lack of similar loss in control plants that underwent the same tissue culture process but no DSB induction. Here too, we do not have direct evidence connecting the CRISPR-Cas-induced DSB and the chromosomal loss, except not finding such events in SpCas9 control plants. However, at least in two cases (plants 1-1 and 5-4), the deletions took place precisely at the DSB site and did not occur in controls, suggesting that these are bona fide DSB induced events rather than being tissue culture-induced.

The analyses of plants 1-1 and 5-4, including genotyping around the break, phenotypic marker analysis, whole-genome sequencing and cytological analyses, supported the hypothesis of the presence of large chromosomal rearrangements, consistent with BFBC (McClintock, 1941). In plants, BFBC has been characterized by McClintock several decades ago (McClintock, 1941). Interestingly, McClintock’s BFBC turned out to be induced by a double-Ds transposon, which might have generated DSBs more challenging for the cell to repair than those made by simple Ac/Ds elements which generate simple excision footprints rather than large-scale chromosomal rearrangements (Weil and Wessler, 1993; English, Harrison and Jones, 1993). Likewise, in this work, one of the loci did not trigger BFBC while the other (on chromosome 11 in an heterochromatic region) did. CRISPR-Cas-induced BFBC has not yet been reported in plants and understanding which loci are repaired in a “clean” manner and which ones undergo a defective repair that generates large-scale genomic rearrangements is very limited.

In mammalian cells, the phenomenon of CRISPR-Cas-induced BFBC has been recently reported, and it is an important concern for the field of gene therapy (Zuccaro et al., 2020; Alanis-Lobato et al., 2021; Leibowitz et al., 2021). There, it is associated with a series of massive rearrangements, including micronuclei formation and translocation of chromosomal segments into new chromosomal locations a phenomenon called chromothripsis (Kwon, Leibowitz, M. and Lee, 2020; Ostapińska, Styka and Lejman, 2022). We report here on a similar syndrome of catastrophic chromosomal rearrangements, namely the occurrence of BFBC, micronuclei, chromosome loss, and translocations, showing that CRISPR-Cas-induced breaks can trigger chromothripsis-like phenomena in plants. The genetic system developed here, with easy-to-monitor phenotypic markers paves the way to study the phenomenon of DSB-induced chromothripsis in plants.

## Materials and Methods

### Plant material

*Solanum lycopersicum* M82 *clv3-1* seeds were a kind gift from the Zachary B Lippman lab (Xu et al., 2015). MT pX11 seeds were a kind gift from the Asaph Aharoni lab (Polturak et al., 2017). Tomato plants were grown in a greenhouse in 5L pots with controlled climate conditions of 26±1°C and a light period of under 12hrs.

### Plasmids

DSB induction was performed by *Streptococcus pyogenes* Cas9 (SpCas9). Arabidopsis optimized SpCas9 was expressed under Parsley Ubiquitin promoter, Ubi4-2, and Pea 3A terminator (Fauser, Schiml, and Puchta, 2014). The gRNAs were expressed under the Arabidopsis U6-26 promoter. A kanamycin resistance gene was expressed under the *Nopaline Synthase* (Nos) promoter and Nos terminator (referred to as Nos:NptII:Nos). Plasmids were cloned using the Golden Braid system (Sarrion-Perdigones et al., 2014).

### Plant transformation

Transformation of tomato plants was done using *Agrobacterium tumefaciens* containing our cloned plasmid. M82 cotyledons transformation was done according to a protocol previously described (Dahan-Meir et al., 2018).

### Genomic DNA extraction

Two or three small tomato leaflets were collected into 1.5ml tubes and ground. Extraction was done according to a protocol previously described (Dahan-Meir et al., 2018), except for genomic DNA for whole-genome sequencing that was extracted using NucleoSpin DNA purification kit (MACHEREY-NAGEL®).

### DNA amplification and sequencing

CRISPR-Cas9 footprints of the parent MT SpCas9 + PSY1 gRNA plant was analyzed using HTS of PCR amplicons (Dahan-Meir et al., 2018). CRISPR-Cas9 DSB regions of the F1 plants [(MT SpCas9 + PSY1 gRNA)/ (M82 pX11 on chromosome 3)], parent M82 *clv3* SpCas9 + gRNA2 plant 2, and F1 plants [(M82 *clv3* SpCas9 + gRNA2)/ (MT pX11 on chromosome 11)], were PCR amplified, and Sanger sequenced. We used the TIDE (Tracking of indels by decomposition) web tool for analysis of the Sanger sequencing CRISPR-Cas9 footprints (Brinkman et al., 2014).

### Inverse PCR

An inverse PCR protocol was used to determine the location of pX11 (Betalain) T-DNA in the M82 and MT lines. Nested primers were designed and used in the inverse PCR reaction (Thomas et al., 1994) (Supplementary Table S1). 300ng of genomic DNA from leaves of M82 and MT pX11 lines were incubated at 37°C overnight with PstI-HF or HindIII-HF (New England BioLabs®) restriction enzymes respectively. Followed by 20 minutes at 65°C for restriction enzymes inactivation. 150ng of the digested fragments were then self-ligated with T4 DNA ligase (New England BioLabs) for two hours at room temperature. The self ligation reaction was then used for the first and second PCR reactions with nested primers, between which the PCR products were cleaned using magnetic beads. The second PCR products were cleaned and sequenced by Sanger sequencing to identify the genomic region flanking the T-DNA left border. Based on the putative genomic integration site primers were designed for amplification and sequencing of the genomic sequences and the LB or RB junctions. An inverse PCR protocol was also used to detect the translocation of chromosome 11 sequence into chromosome 9 in the 5-4 green F1 plant. Genomic DNA was digested with PstI restriction enzyme. Nested primers were designed and used in the inverse PCR reaction (Thomas et al., 1994) (Supplementary Table S6). The rest of the protocol is as above. We sequenced the PCR product by Sanger sequencing to identify the genomic region of the translocation downstream junction. Additional PCR primers were designed for verification and Sanger sequencing of the chromosome 11 into chromosome 9 translocation junctions (Supplementary Table S7)

### Plant regeneration

F1 plants were sterilized and sown in boxes containing Nitsch growth medium, and once grown, their leaves were dissected to small ∼0.5 cm^2^ pieces. They were then transferred to the selection I medium, as described in the plant transformation section above, but without kanamycin. The leaves were then gradually transferred to plates containing selection mediums II and III, and then to boxes with rooting medium and to 5L pots in the greenhouse.

### SpCas9 + gRNA transgenic plant generation

The gRNAs were cloned into a Golden braid vector (Sarrion-Perdigones et al., 2014). PSY1 gRNA was cloned into p3Ω1 nos:nptII:nos ubi:SpCas9 U626:PSY1 gRNA. gRNA2 was cloned into p3Ω1 nos:nptII:nos ubi:SpCas9 U626:gRNA2.

The MT SpCas9 + PSY1 gRNA plant #10 was previously selected (Dahan-Meir et al., 2018), and was used for crosses with M82 pX11 plants. The heterozygous (50% -GCT/ 50% -G) footprint in the gRNA recognition sequence in this plant created the conditions for an allele-specific assay, in which a DSB will occur only in the M82 parent (Supplementary Table S2).

M82 *clv3* cotyledons were transformed with the gRNA2 cassette, regenerated rooted plants were transferred successfully to the greenhouse, and transformant plant tissue was collected for DNA extraction and sent to Sanger sequencing of the target area. Presence of SpCas9 was confirmed in all plants through PCR amplification with primers from within the Cas9 sequence (Supplementary Table S3). Different NHEJ repair footprints were observed, and one specific line (M82 *clv3* SpCas9 + gRNA2 plant #2) was chosen for crosses with MT pX11 plants. The homozygous (100% +T) footprint in the gRNA recognition sequence in this plant created the conditions for an allele-specific assay, in which a DSB will occur only in the MT parent (Supplementary Table S2).

### SNP analysis by Sanger sequencing

To analyze SNPs between M82 and MT at the DSB target sites, and from both sides of the chromosome 3 PSY1 gRNA or chromosome 11 gRNA2 targets, we performed PCRs of each selected SNP region, on genomic DNA. The PCR products were then sequenced by the Sanger method. The details of the PCR primers and their genomic location are in Supplementary Table S5.

### Whole genome sequencing and analysis

DNA was purified from leaves of F1, and F2 plants using a DNA purification kit (MACHEREY-NAGEL®) and then 300 ng sheared by sonication (Bioruptor ®, Diagenode) to 200–500 bp. A total of 10 ng of fragmented DNA per plant was used for libraries preparation as described in (Ben Shlush et al., 2020). High-throughput sequencing was performed at the Life Sciences Core facilities unit at the Weizmann Institute of Science with the Illumina NovaSeq 6000, 150 bp paired-end reads. The coverage of the various genomes was x5-x31 (Supplementary Table S8). The whole genome sequencing reads of the selected tomato plants were aligned to the SL4.0 tomato genome version (Sol Genomic Network), as in (Ben Shlush et al., 2020). The reads were viewed and further analyzed using the IGV browser (Robinson et al., 2011).

### Coverage analysis

Average coverage was computed using GATK version 3.7. The genome was divided in windows of 250kb for genome-wide analyses, and 5kb for more detailed inspections of the expected DNA double-strand break site. Dosage was estimated by normalizing the observed number of reads per window by the mean number of reads, stratified by GC content to account for differences in coverage due to GC content bias. GC content was computed with bedtools nuc version 2.26 across windows. Windows were then subdivided in deciles of GC content, and the mean number of reads was estimated for each decile, and then used for normalization. The presence of chromosomal rearrangements on each side of the Cas9 target site was tested comparing with a likelihood ratio test models in which no rearrangements, a full deletion of both chromosomes, a single chromosomal arm deletion or a duplication, occurred after the break assuming a dosage of 2x, 0.01x,1x or 3x, respectively, and considering 1Mbp before and after the break. Rearrangements supported by a p-value lower than 0.001 were considered as true rearrangements. The R code used for plotting and testing of the rearrangements can be found at github.com/fabrimafe/CRISPRcoverage.

### Imaging of meiocytes

Tomato flower buds were fixed in ethanol: acetic acid 3:1 for at least 1 hour, then kept in 70% ethanol at 4^◦^C. Anthers of flower buds were dissected and stained in 50ng/µl DAPI in ProLong Gold Antifade Mountant. For imaging, we used Nikon Eclipse Ti microscope, and x100 Nikon N plan Apo lambda 100x/1.45 oil OFN25 DIC N2 lens. Images were transferred into the DeltaVision (Applied Precision) format, deconvolved, and examined in 3D. The images presented in Figure 8, are flat projections of the 3D images.

## Author Contributions

Conceptualization, A.S. and A.A.L.; methodology, A.S., O.G., C.M.-B., T.D-M., S.F.-H., Z.A., and W.P.P.; software, F.M.; validation, A.S., F.M., C.M.-B., and W.P.P.; writing original draft preparation, O.G., A.S., and A.A.L.; writing review and editing, A.S., F.M., O.G., C.M.-B., T.D.-M., S.F.-H., Z.A., W.P.P., and A.A.L.; supervision, A.A.L.; project administration, A.S. and A.A.L.; funding acquisition, A.A.L. All authors have read and agreed to the published version of the manuscript.

## Acknowledgments

The authors would like to thank Prof. Asaph Aharoni for the MT pX11 plant materials; Prof. Zachary Lippman for the *clv3-1* mutant plant materials; the Weizmann Institute greenhouse staff for the plants’ maintenance; Dr. Gil Feiguelman for guidance with microscopy and all the Levy lab members for fruitful discussions and assistance during the course of this study.

## Supplementary Materials

**Supplementary Figure S1.**
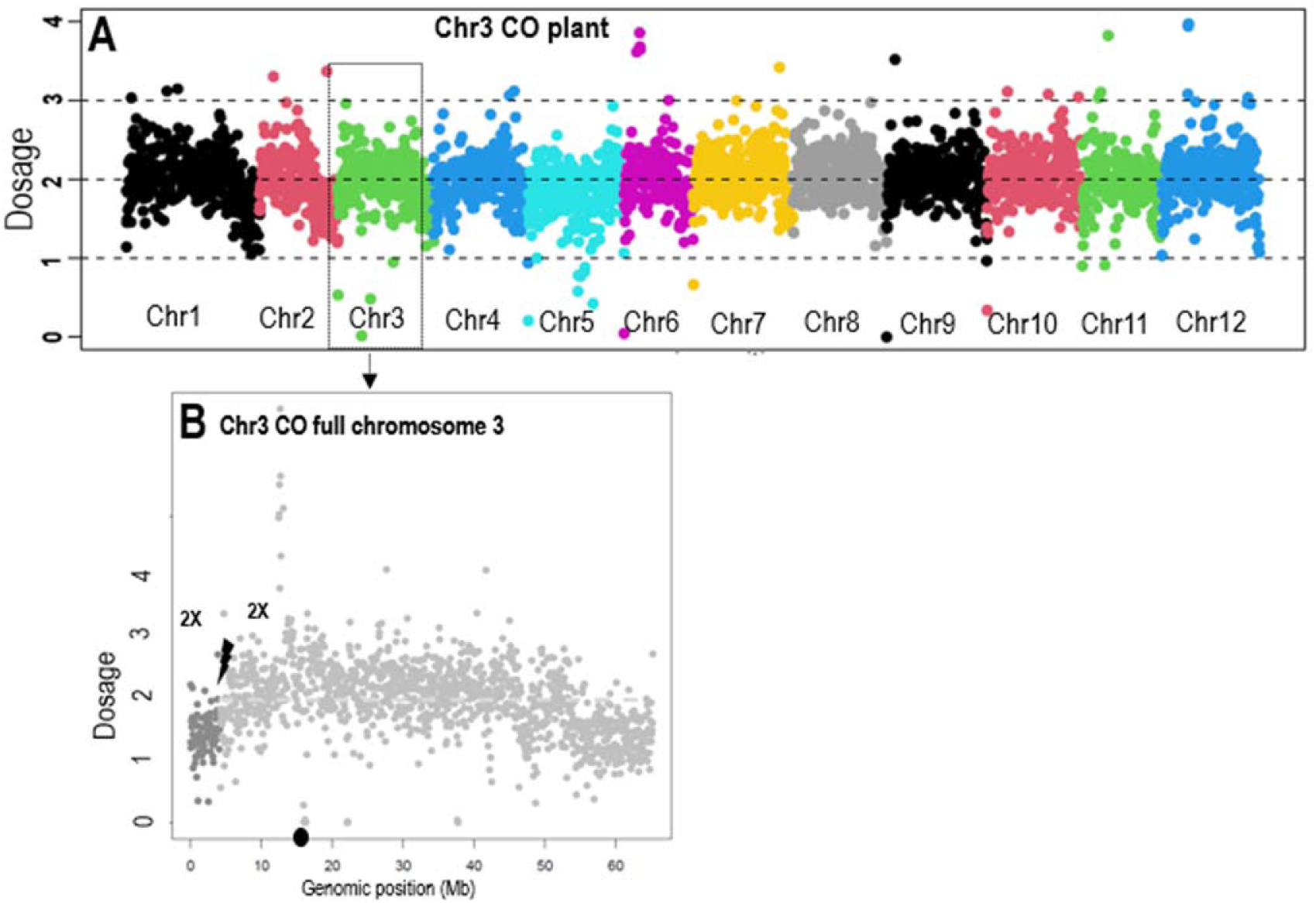
WGS Coverage analyses show no dosage changes at the induced DSB site of chromosome 3 CO event. Average coverage of WGS reads per plant was determined as 2X diploid dosage basis. A: Dosage for each of the 12 chromosomes, is presented with each chromosome shown in a different color. B: Dark grey dots are dosage bins from genomic position 1 up to the DSB site, and light grey dots are dosage bins from the DSB site to the end of chromosome 3. Chromosome 3 CO plant whole chromosome 3 dosage shows similar ∼2X dosage in both sides of the DSB site. This indicates that the genotype transition from both sides of the DSB is not due to loss of the region from the DSB site to the telomere in one of the chromosomes. Chr-chromosome. Black lightning bolt – DSB site. Black dot on Genomic position (Mb) axis – centromere.

**Supplementary Figure S2.**
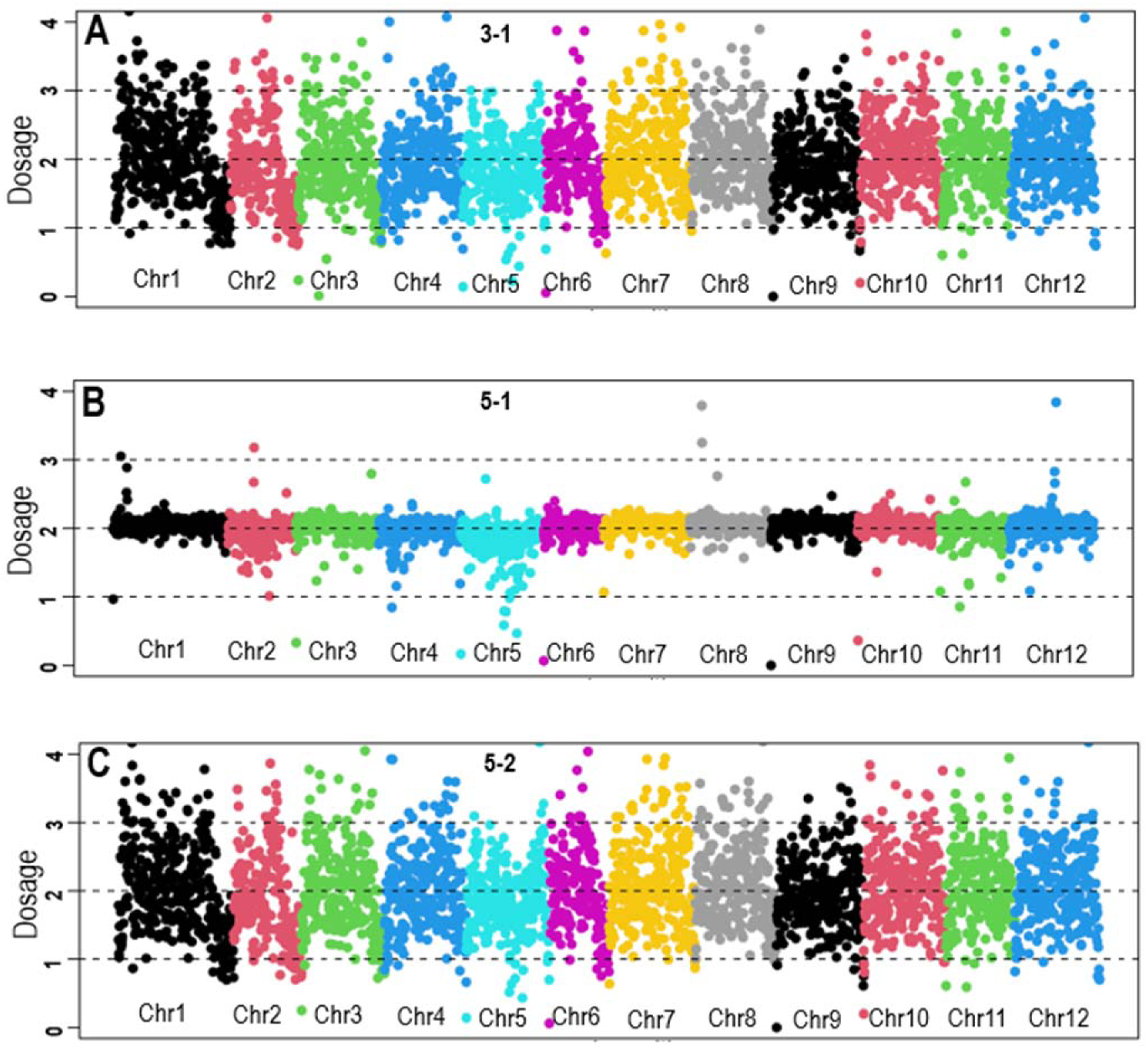
WGS Coverage analyses show no dosage changes at the induced DSB site of chromosome 11 T-DNA loss events. Average coverage of WGS reads per plant was determined as 2X diploid dosage basis. A-C: Coverage for each of the 12 chromosomes per plant, is presented with each chromosome shown in a different color. Panel A: plant 3-1. Panel B: plant 5-1. Panel C: plant 5-2. The dosage in these three plants is 2X in all chromosomes. Chr-chromosome.

**Supplementary Figure S3.**
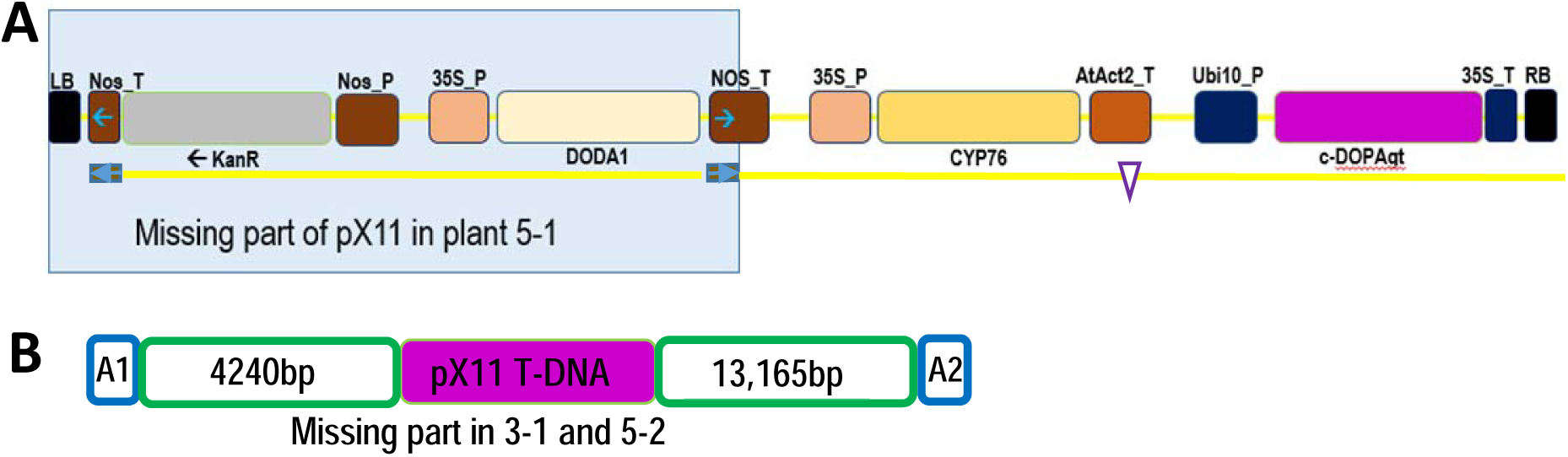
Loss of Betalain expression cassette T-DNA (pX11) in plants 5-1, 3-1 and 5-2. This type of deletion events of plant 5-1, 3-1 and 5-2 was detected only in a plant with Cas9 and where a DSB was induced and not in control plants. The distance between the DSB and the end of the T-DNA integration sites is 180,895bp. F2 progeny of each of these plants that were homozygous to MT SNPS in the regions flanking the gRNA recognition site and the pX11 T-DNA insertion site, were analyzed by WGS. In the deleted region there were no reads. A: In plant 5-1 a part of the pX11 T-DNA was lost. The region with a blue box behind it is the part missing in plant 5-1. The brown boxes with blue arrows in them are the NOS terminator repeated sequences in inverted orientations. These sequences could anneal and generate a loop of the sequence between them. The empty purple triangle indicates the pX11 cassette’s sequences that are present in plant 5-1 but do not give the Betalain color. KanR: kanamycin resistance gene; DODA1: B. vulgaris DOPA 4,5-dioxygenase; CYP76: B. vulgaris cytochrome P450; cDOPAgt: M. jalapa cyclo-DOPA-5-O-glucosyltransferase;Nos P/T: nopaline synthase promoter/terminator; 35S P/T: CaMV 35S promoter/terminator; AtAct2_T:Arabidopsis actin 2 terminator; Ubi10_P: Arabidopsis ubiquitin 10 promoter. B: In plants 3-1 and 5-2, two independent F1 regenerated green plants, the same region of MT chromosome 11 was deleted. A1 and A2 are A rich repeats flanking the deleted region.

**Supplementary Figure S4.**
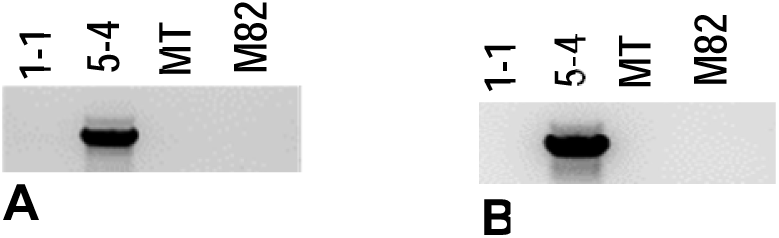
PCR amplification of chromosome 11 translocation into chromosome 9 junctions from plant 5-4. Two PCR primers sets were used for amplification of the chromosome 11 translocation into chromosome 9 junctions (Supplementary Table S6.). In both cases, a PCR product specific to plant 5-4 was amplified. A: Amplification of the downstream junction with primers gRNA_pair2_far_F and CHR9_insert_CH11_GR2_R. B: Amplification of the upstream junction with primers CH9_trans_US_F2 and gRNA_pair2_far_F.

**Supplementary Figure S5.**
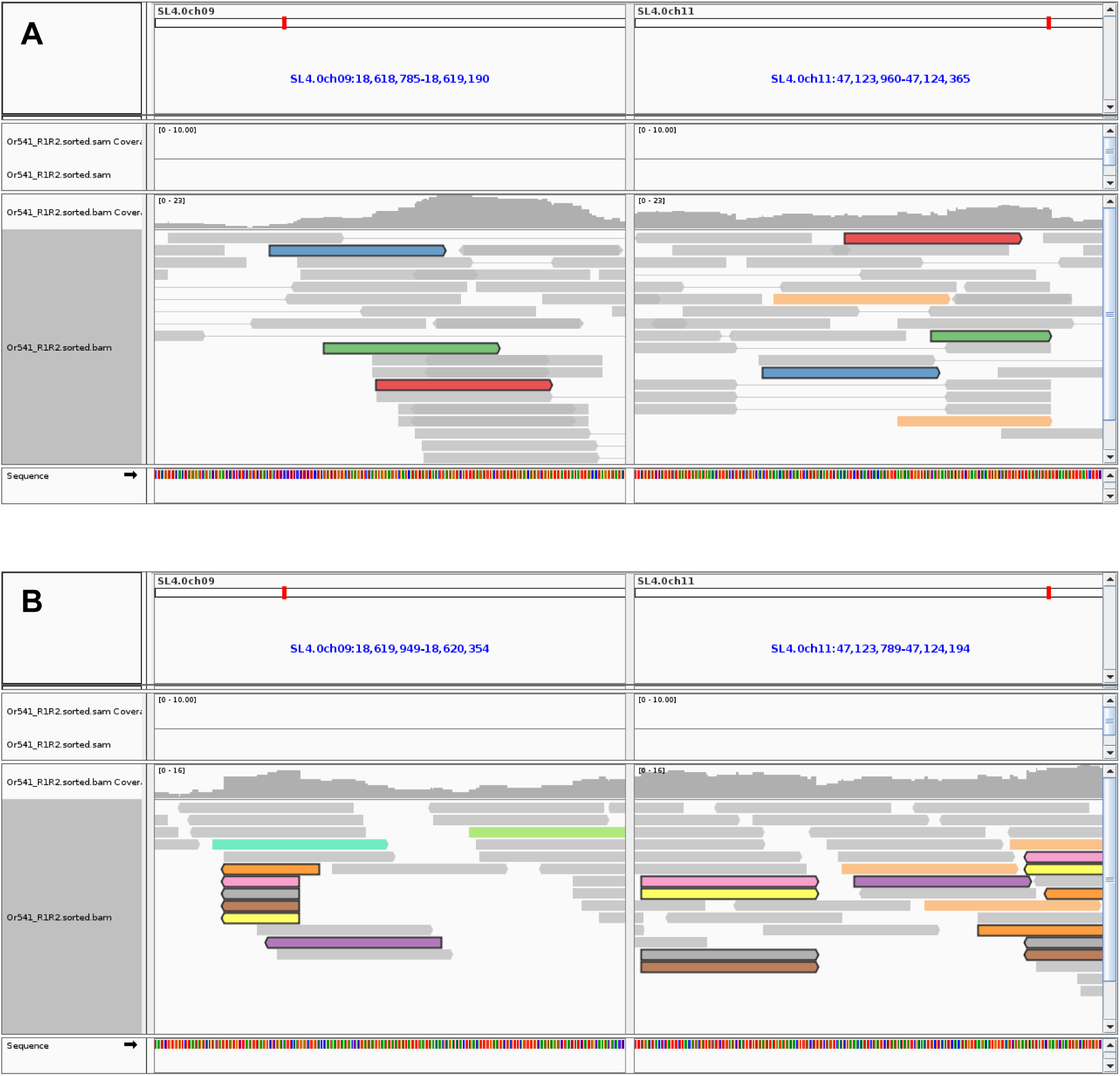
Illumina reads of chromosome 11 translocation into chromosome 9 junctions. IGV presentation of the Illumina reads of chromosome 11 translocation into chromosome 9 junctions. Reads pairs with black lining and the same color indicate either pairs in which the two reads map to the two different chromosomes, or split reads in which a single read out of the two span the putative translocation site and thus map to two different chromosomes. A: Reads of the upstream junction of chromosome 9 and chromosome 11. In this case, no individual reads span the translocation site, while three read pairs are formed by individual reads mapping to chromosome 9 and chromosome 11. B: Reads of the downstream junction of chromosome 9 and chromosome 11. Here one read pair is formed by two individual reads falling entirely on chromosome 9 or chromosome 11 (purple), while five out of six reads span the translocation junction between the two chromosomes, thus appearing truncated and on both sides of the panel, i.e. both on chromosome 9 and 11. These reads therefore represent the transition from chromosome 9 to chromosome 11 sequences.

**Supplementary Table S1.**
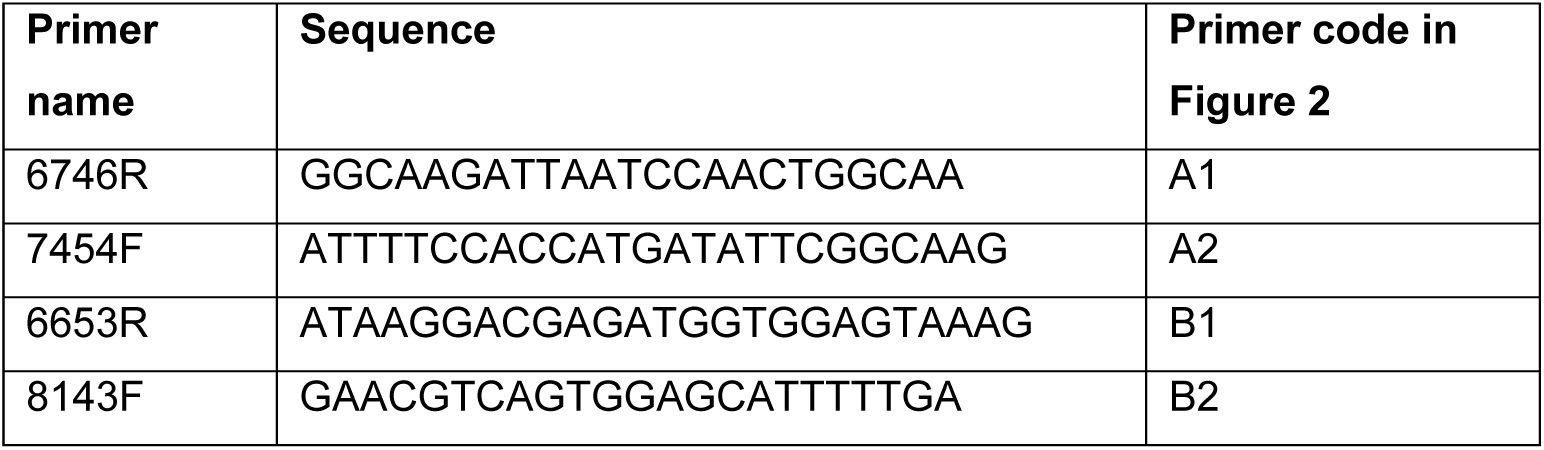
LB primers for inverse PCR. Nested primers used for the inverse PCR of the T-DNA left border (LB). The first pair, 6746R + 7454F, is the inner pair equivalent to A1+A2 in figure 2. The second pair, 6653R + 8143F, is the outer pair equivalent to B1+B2 in figure 2.

**Supplementary Table S2.**
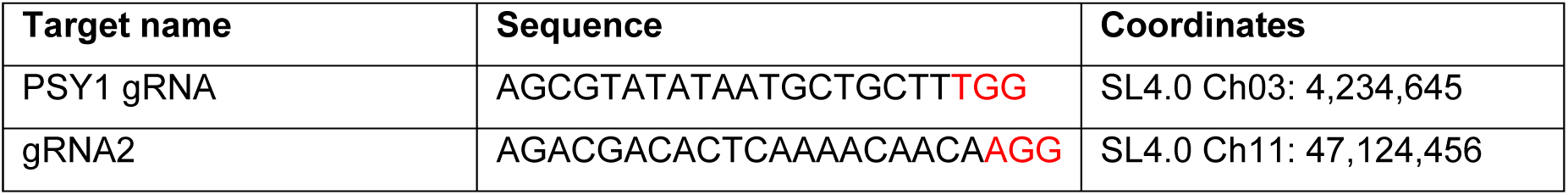
Cas9 DSB Targets on chromosome 3 and chromosome 11. The sequence and coordinates for both targets. PAM sequence (NGG) in each target is marked in red.

**Supplementary Table S3.**
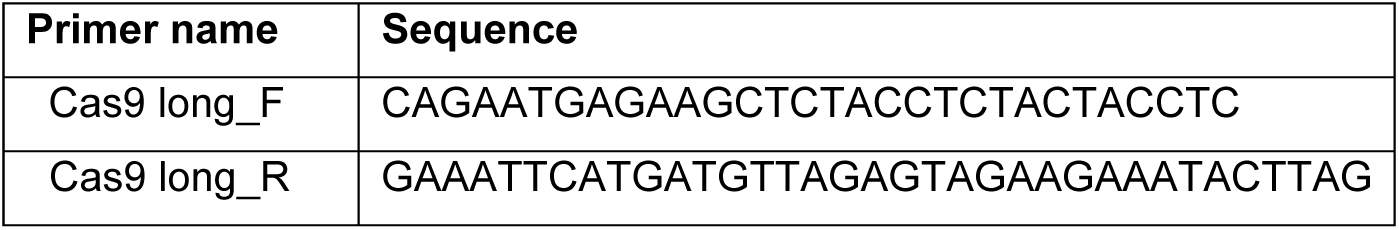
PCR Primers used for SpCas9 positive plants screening and selection. SpCas9 primers for verification of Cas9 T-DNA cassette presence in transgenic plants.

**Supplementary Table S4.** Silencing and LOH events frequencies. Number of silencing and LOH events in chromosome 3 or chromosome 11. Plants grown in the greenhouse or cut and regenerated in tissue culture. Control plants with SpCas9 only, and treatment plants with SpCas9 + gRNA.

|  | Chromosome 3 PSY1 gRNA |  |  |  | Chromosome 11 gRNA2 |  |  |  |
| --- | --- | --- | --- | --- | --- | --- | --- | --- |
|  | M82/MT F1<br>in greenhouse |  | M82/MT F1<br>in tissue culture |  | M82/MT F1<br>in greenhouse |  | M82/MT F1<br>in tissue culture |  |
|  | SpCas9 | SpCas9<br>+gRNA | SpCas9 | SpCas9<br>+gRNA | SpCas9 | SpCas9<br>+gRNA | SpCas9 | SpCas9<br>+gRNA |
| Number of plants | 10 | 10 | 10 | 10 | 10 | 10 | 12 | 9 |
| Number of silencing events | 0 | 0 | 1 | 2 | 0 | 0 | 1 | 1 |
| Number of LOH T-DNA loss events | 0 | 0 | 0 | 0 | 0 | 0 | 0 | 3 |
| Number of LOH chromosome arm loss | 0 | 0 | 0 | 0 | 0 | 0 | 0 | 2 |
| Number of LOH whole chromosome loss | 0 | 0 | 0 | 0 | 0 | 0 | 0 | 1 |
| Number of LOH crossover | 0 | 1 | 0 | 0 | 0 | 0 | 0 | 0 |

**Supplementary Table S5.**
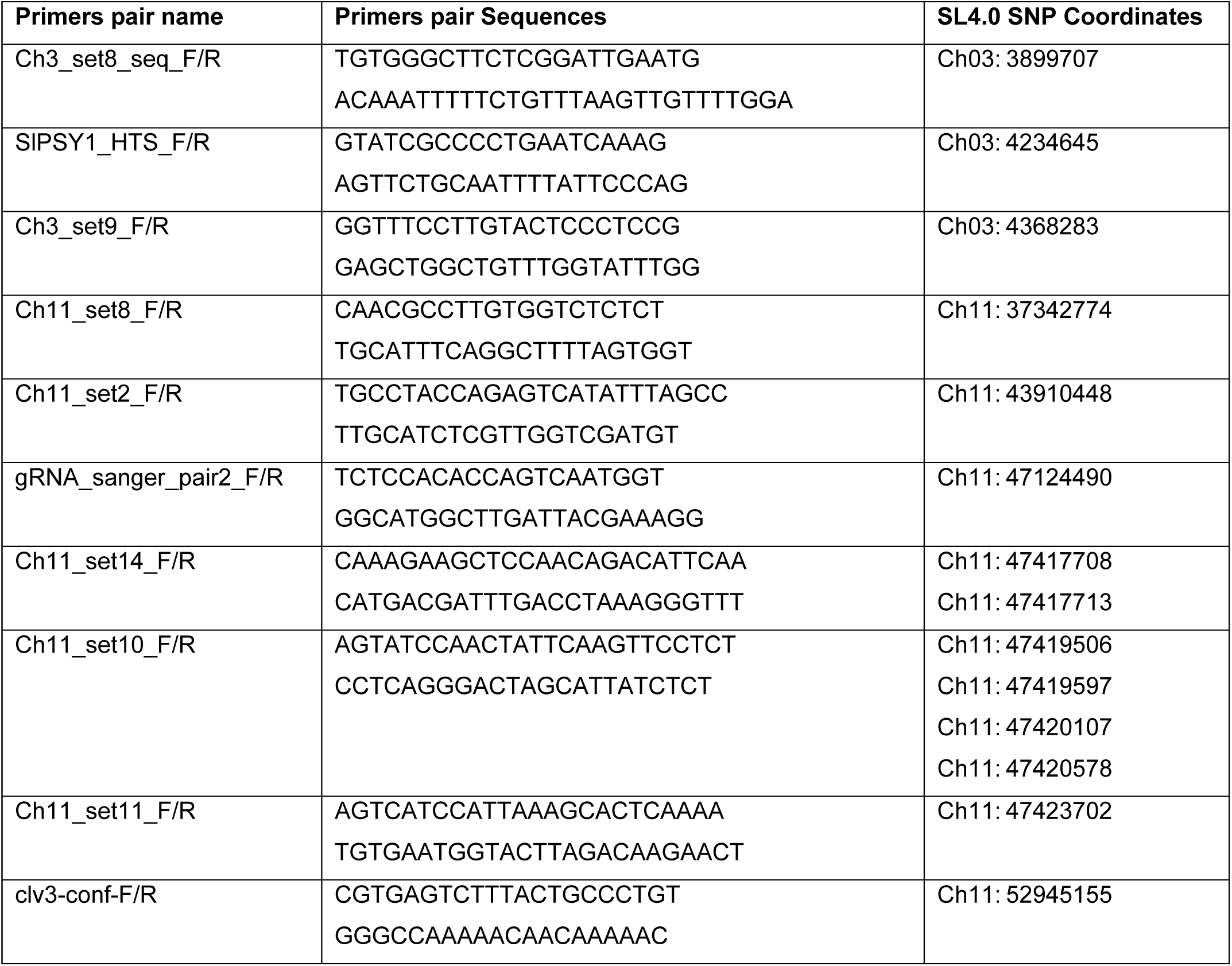
Primers used for sequencing of SNPs in F1, F2 and F3 plants. Each primers pair was used for PCR amplification of the SNP region. One or both primers of each set were used for Sanger sequencing of the PCR amplicon.

**Supplementary Table S6.**
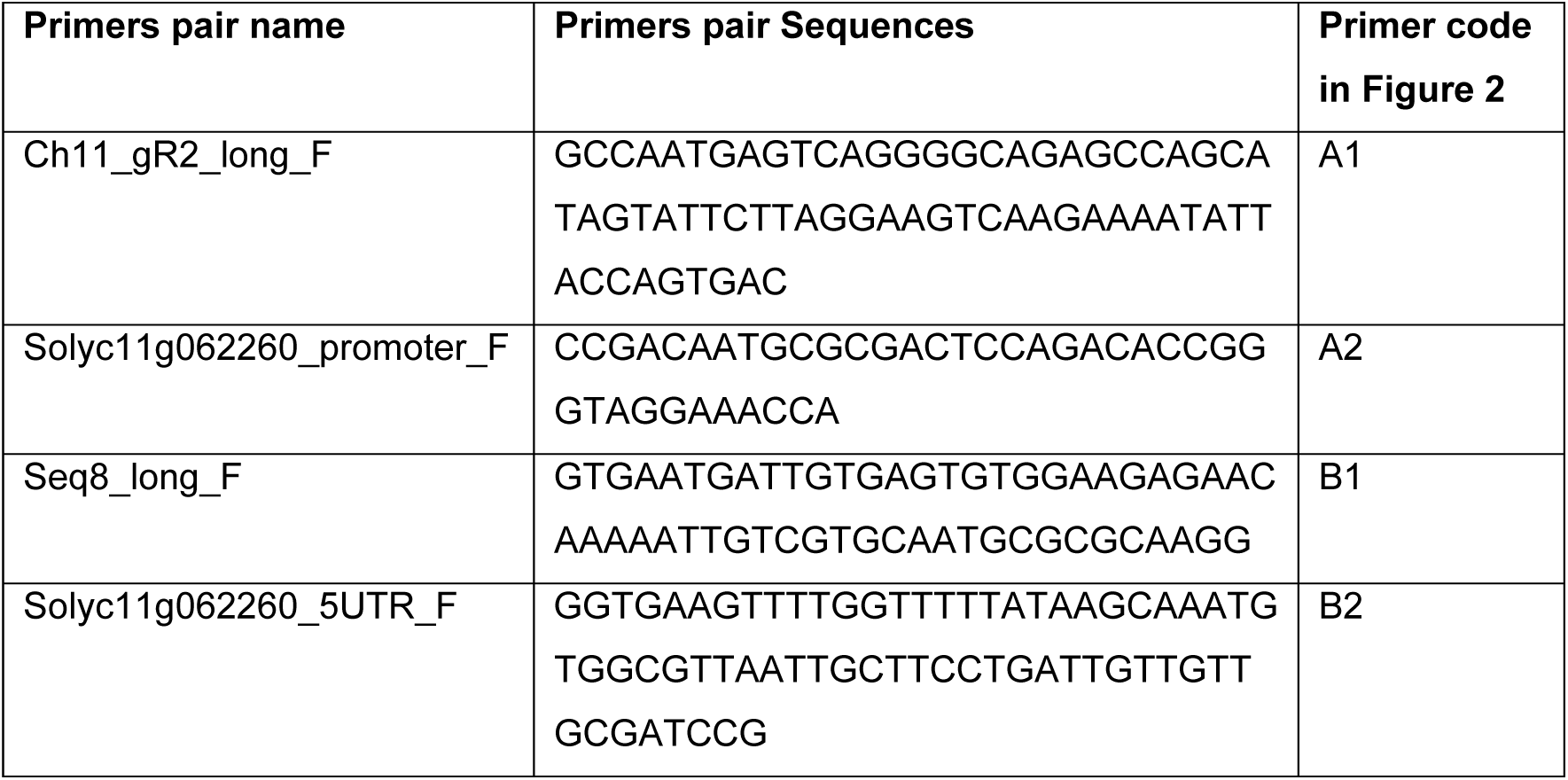
Primers used for inverse PCR and detection of plant 5-4 chromosome 11 into chromosome 9 translocation junctions. Nested primers used for the inverse PCR and detection of plant 5-4 chromosome 11 into chromosome 9 translocation. The first pair, Ch11_gR2_long_F + Solyc11g062260_promoter_F, is the inner pair equivalent to A1+A2 in figure 2. The second pair, Seq8_long_F + Solyc11g062260_5UTR_F, is the outer pair equivalent to B1+B2 in figure 2.

**Supplementary Table S7.**
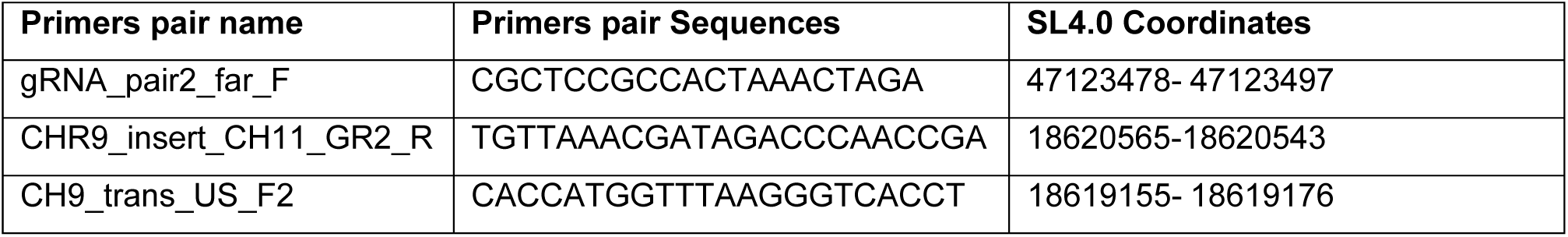
Primers used for sequencing of plant 5-4 chromosome 11 into chromosome 9 translocation junctions. Each primers pair was used for PCR amplification of the junction region. One or both primers of each set were used for Sanger sequencing of the PCR amplicon. gRNA_pair2_far_F is on chromosome 11 side of both junctions, and was paired with each of the chromosome 9 primers.

**Supplementary Table S8.** Whole genome sequencing (WGS) coverage. For WGS we used Illumina NovaSeq 6000, with 150 bp paired-end reads. Coverage was calculated as read length (300bp) x number of reads / haploid genome length (Tomato SL4.0 = 782,520,133 bp). Chr-chromosome, MT-Micro-Tom

| Plant | Plant number | Number of reads<br>(x million) | Coverage | Library Complexity<br>(% of duplicates) |
| --- | --- | --- | --- | --- |
| Chr 3 CO | F2 124 | 33.2 | X12 | 14% |
| Chr11 T-DNA loss | 3-1 (F2 22) | 13.7 | X5 | 9% |
| Chr11 T-DNA partial loss (LB, KanR, DODA) | 5-1(F2 10) | 81.4 | X31 | 7% |
| Chr11 T-DNA loss | 5-2 ( F2 24) | 24.6 | X9 | 11% |
| Chr11 arm loss | 1-1 | 17.9 | X6 | 5% |
| Chr11 arm loss | 5-4 | 28.9 | X11 | 12% |
| Chr11 loss | 9-1 | 43.6 | X16 | 15% |
| MT |  | 65.5 | X25 |  |
| M82 |  | 55 | X21 | 18% |

